# Fyn-mediated phosphorylation of Menin disrupts telomere maintenance in stem cells

**DOI:** 10.1101/2023.10.04.560876

**Authors:** Souren Paul, Preston M. McCourt, Le Thi My Le, Joohyun Ryu, Wioletta Czaja, Ann M. Bode, Rafael Contreras-Galindo, Zigang Dong

**Author notes:** Correspondence to: Souren Paul, The Hormel Institute University of Minnesota, Austin, MN, 55912, USA, Rafael Contreras-Galindo Department of Genetics, University of Alabama, Birmingham, AL 35294, USA, Zigang Dong, Zhengzhou University, Zhengzhou, Henan, 450001, China.

## Abstract

Telomeres protect chromosome ends and determine the replication potential of dividing cells. The canonical telomere sequence TTAGGG is synthesized by telomerase holoenzyme, which maintains telomere length in proliferative stem cells. Although the core components of telomerase are well-defined, mechanisms of telomerase regulation are still under investigation. We report a novel role for the Src family kinase Fyn, which disrupts telomere maintenance in stem cells by phosphorylating the scaffold protein Menin. We found that Fyn knockdown prevented telomere erosion in human and mouse stem cells, validating the results with four telomere measurement techniques. We show that Fyn phosphorylates Menin at tyrosine 603 (Y603), which increases Menin’s SUMO1 modification, C-terminal stability, and importantly, its association with the telomerase RNA component (TR). Using mass spectrometry, immunoprecipitation, and immunofluorescence experiments we found that SUMO1-Menin decreases TR’s association with telomerase subunit Dyskerin, suggesting that Fyn’s phosphorylation of Menin induces telomerase subunit mislocalization and may compromise telomerase function at telomeres. Importantly, we find that Fyn inhibition reduces accelerated telomere shortening in human iPSCs harboring mutations for dyskeratosis congenita.

## Introduction

Telomeres, regions of TTAGGG repeats at the ends of the chromosomes, shrink in somatic cells during cell replication^1^. Telomere shortening can be counteracted by telomerase holoenzyme, a ribonucleoprotein that maintains telomere length by synthesizing telomeric DNA^2^. Although telomerase activity is greatly reduced in somatic cells, it is upregulated in stem cells that undergo rapid expansion^3^. In telomere biology disorders (TBD) such as dyskeratosis congenita, telomere maintenance is defective, which often causes bone marrow failure due to accelerated telomere shortening in hematopoietic stem cells ^4^. Aberrant telomere maintenance in dyskeratosis congenita is most frequently a consequence of mutations in telomerase holoenzyme or shelterin complex components^5^. Currently, there is no specific treatment for TBD and transplantation comes with great risk and poses a number of challenges^6^.

Endogenous telomere maintenance is modulated by many methods, including transcriptional regulation of the telomerase RNA component (TR) and telomerase reverse transcriptase (TERT), post-translational regulation of TERT, and telomerase recruitment and assembly^7^. Little is known about the role that kinases play in telomere maintenance, although previous studies showed that ATM and ATR are positive regulators of telomere length^8, 9^.

Fyn is a non-receptor Src family tyrosine protein kinase that phosphorylates multiple substrates and regulates many critical biological processes, including cell growth and survival^10,11^. Previous studies showed that Src family kinase inhibition prevents stem cell differentiation^12,13^. Here, we describe a critical role for Fyn, which negatively regulates telomere length in mammalian stem cells. We determined that genetic or pharmacological suppression of Fyn stabilizes telomere length in mouse and human stem cells. We also found that post-translational modifications of Menin, a novel Fyn phosphorylation substrate, play crucial roles in Fyn-mediated telomere erosion. Named for its role in multiple endocrine neoplasia type 1 (MEN1) syndrome, Menin is a scaffold protein that regulates gene expression, interacts with multiple signaling pathways, and can act as a cancer suppressor or promoter depending on the context^14^. A previous study found that Menin localizes to telomeres in meiotic cells but does not directly interfere with telomerase activity in carcinoid cells^15^. Here we show that Fyn promotes Menin localization to telomeres in mouse embryonic stem cells (mESCs) and that Fyn does not modulate telomerase activity using *in vitro* assays. We also show that Menin is phosphorylated by Fyn at Y603 promoting phosphorylation-dependent SUMO1 modification at its C-terminal region. Importantly, we demonstrate that SUMO1-modified Menin has greater stability and increased affinity for TR, which disrupts TR’s association with the telomerase subunit Dyskerin likely after telomerase assembly. These results suggest that SUMO1-modified Menin induces telomerase subunit mislocalization and may compromise telomerase activity. Thus, Fyn may act as a negative regulator of telomere maintenance by phosphorylating and promoting stable SUMO1-modification of Menin. We further demonstrate that Fyn knockout prevents accelerated telomere erosion in DC patient iPSCs that harbour a mutation in DKC1. Our findings indicate that Fyn inhibitors may be a viable therapeutic intervention for patients with TBD.

## Results

### Fyn inhibition slows telomere erosion in mice and stem cells

We previously found that Fyn^−/−^ mice produced larger and greater number of tumors after induction of skin carcinogenesis^10^. Other studies have focused on the role of Fyn in stem cell differentiation, finding that Src family kinase inhibition prevented stem cell differentiation^12,13^. Given that telomere maintenance is a critical component of both cancer and stem cell proliferation, we were curious to see how the absence of Fyn affected telomere length in Fyn^−/−^ mouse tissue and stem cells. In a pilot study, we found that 43-week-old Fyn^−/−^ female mice (n=4) exhibited greater telomere length in colon, and skin tissues compared to WT mice, while centromere length was unchanged (Supplementary Figs.1-2). We previously demonstrated that centromere size does not change with cell passaging^16^. Therefore, we used centromere Q-FISH as a baseline control. Representative Q-FISH images (Supplementary Figs.1, 2) and frequency distribution of mean fluorescence with detailed statistics (Supplementary Figs. 1B, C; 2B, C) from these images show longer telomere length in Fyn^−/−^ mouse colon, and skin tissues compared to WT. Statistical analysis showed a significantly higher telomere mean fluorescence (Supplementary Figs. 1D, 2D), but no significant difference was noted when centromere mean fluorescence was compared (Supplementary Figs. 1E, 2E) between WT and Fyn^−/−^ mouse tissues. The telomere/centromere mean fluorescence ratio was also significantly higher in Fyn^−/−^ mouse compared to WT (Supplementary Figs. 1F, 2F) in colon, and skin. As previous reports showed that skin and gastrointestinal tract tissues are excellent sources of stem cells^17,18^, we hypothesized that Fyn may negatively regulate telomere length in stem cells.

We tested this hypothesis in mESCs obtained from WT and Fyn^−/−^ blastocysts (Fig. 1A, C, and Supplementary Figs. 3A-G). Fyn knockout was confirmed by western blot (Fig. 1C). Q-FISH analysis showed that telomeres in Fyn^−/−^ stem cells are indeed longer than telomeres in WT stem cell metaphase spreads (Fig. 1A) or interphase nuclei (Supplementary Figs. 3E-F) at passage 12. We also performed Fyn knockdown in ES-E14TG2a (E14) mESCs using sh-RNA and confirmed knockdown by western blot (Fig. 1D). Sh-Fyn cells showed greater telomere length compared to sh-Mock cells (Fig. 1B, and Supplementary Figs. 3E, 3G) at passage 18.

**Fig. 1.**
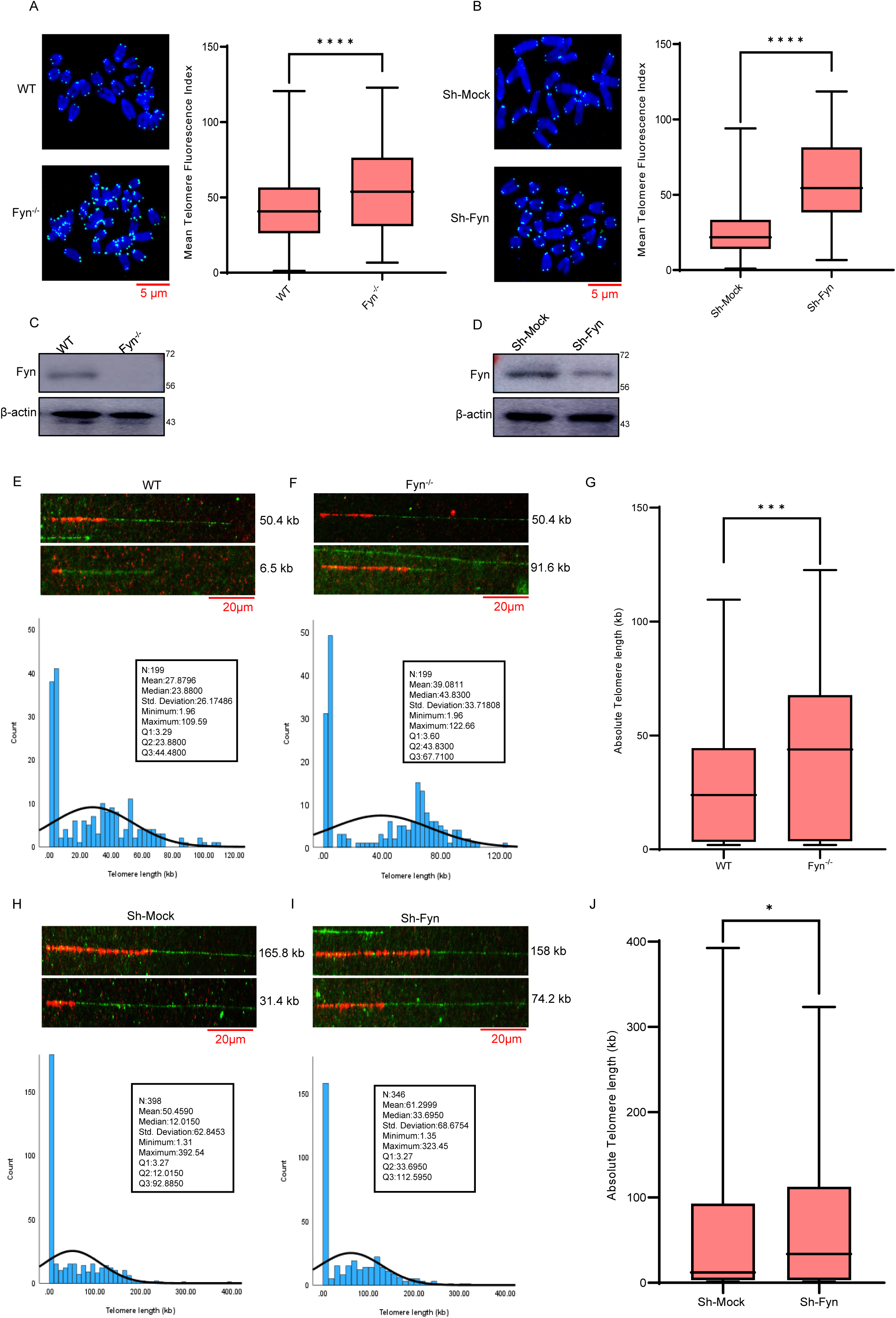
Fyn inhibition slows telomere erosion in mESCs and tissue. Representative images of Q-FISH telomere analysis in metaphase chromosomes of mESCs isolated from (A) WT and Fyn^−/−^ mice (passage 12) and (B) stable sh-Mock and sh-Fyn E14 cells (passage 18). Differences in the Mean Telomere Fluorescent Index, calculated from 400 telomeres in 20 metaphase spreads, are shown at the right of each figure. Scale bar is 5 µm. Western blotting of Fyn in (C) WT and Fyn^−/−^ mESCs and (D) sh-Mock and sh-Fyn E14 cells. β-actin served as a loading control. Protein molecular weights are shown at the right. Representative images of telomere analysis using Molecular Combing Assays from genomic DNA of mESCs isolated from WT (A) and Fyn^−/−^ (B) mice (passage 12) as well as sh-Mock (D) and sh-Fyn (E) mESCs (passage 12). The scale bar is 20 µm. A detailed statistical analysis and frequency of distribution of telomere sizes in chromatin fibers are shown below sample images. The telomere size was quantified using a telomere probe. Q1, quartile 1; Q2, quartile 2 and Q3, quartile 3. Differences in Absolute Telomere length in kb between WT and Fyn^−/−^ mESCs (C) and between sh Mock and sh Fyn mESCs (F) are shown. Data are presented as means ± SD from a two-tailed t-test (* ^=^ *p* < 0.1, *** = *p*LJ<LJ0.001, **** = *p*LJ<LJ0.0001). A probability of *p* < 0.05 was considered statistically significant.

We used qRT-PCR and Flow-FISH to further confirm that Fyn^−/−^ ES cells and Fyn knockdown E14 cells have greater telomere length (Supplementary Figs. 4A-D). The mean telomere fluorescence value was higher in Fyn-deficient ES cells than WT ES cells or mock-infected cells as determined by Flow-FISH (Supplementary Figs. 4A and 4C). A significantly higher telomere to single copy gene ratio (T/S) was observed in Fyn^−/−^ ES cells (Supplementary Fig. 4B) and Fyn knockdown E14 cells (Supplementary Fig. 4D) compared to WT ES cells and sh-mock cells, respectively. Notably, stable overexpression of the *Fyn* gene in sh-Fyn E14 cells (Supplementary Figs. 4E-4H) significantly reduced telomere FISH signals as compared to sh-Fyn cells (Supplementary Figs. 4E, 4H), suggesting that the level of active Fyn imposes a negative impact on telomeres.

We also investigated whether Fyn^−/−^ mice had longer telomeres than WT mice at old age. Genotyping, IHC analysis and quantification of IHC confirmed Fyn knockout in colon and skin tissues of 97-week-old Fyn^−/−^ mice (Supplementary Figs. 5A-F). Q-FISH analysis showed that telomere length in Fyn^−/−^ mice was significantly greater in both male and female mice compared to WT mice (Supplementary Figs. 6A-F).

Moreover, Telomere Molecular Combing Assay (TMCA) results strengthen our observation that Fyn inhibition promotes telomere maintenance in stem cells (Fig. 1E-J). The TMCA shows that Fyn^−/−^ mESCs DNA fibers have greater telomere length compared to WT mESCs DNA fibers (Fig. 1E, F, G). Detailed statistical analysis from the telomere signals shows that mean and median telomere sizes were 39.0811 kb and 43.8300 kb, respectively, in Fyn^−/−^ mESCs compared to WT mESCs, which shows that mean and median telomere sizes were 27.8796 kb and 23.8800 kb, respectively (Fig. 1E, F). A comparative statistical analysis shows significantly longer absolute telomere length in Fyn^−/−^ mESCs compared to WT mESCs (Fig. 1G). Similarly, Sh-Fyn stable E14 stem cells show mean and median telomere sizes of 61.2999 kb and 33.6950 kb, respectively, compared to Sh-Mock, which shows mean and median telomere sizes of 50.4590 kb and 12.0150 kb, respectively (Fig. 1H, I). Comparative statistical analysis also shows significantly greater absolute telomere length in Sh-Fyn cells compared to Sh-Mock cells (Fig. 1J). Taken together, these results indicate that Fyn is a negative regulator of mammalian telomere length in stem cells.

### Fyn does not inhibit telomerase activity *in vitro*

To elucidate the molecular mechanism of Fyn-mediated telomere erosion, we measured telomerase reverse transcriptase (TERT) expression and found no difference between Fyn^−/−^ and WT stem cells or between Fyn knockdown and sh-mock stem cells (Supplementary Fig. 7A-B). Therefore, the greater telomere length found in Fyn^−/−^ and Fyn knockdown stem cells does not appear to be the result of a change in TERT expression. To evaluate whether Fyn directly interferes with telomerase activity, we performed telomere repeat amplification protocol (TRAP)^19^. There was no significant difference in telomerase activity between Fyn^−/−^ and WT mESCs (Supplementary Fig.7C), Sh-Fyn and Sh-Mock E14 cells (Supplementary Fig. 7D), or colon lysates of Fyn^+/+^ and Fyn^−/−^ mice (Supplementary Fig. 7E) and in Sh-Mock vs Sh-Fyn HEK293T cells (Supplementary Fig. 7F). The TRAP assay is helpful for measuring telomerase activity, but cannot detect improper localization of telomerase subunits^20, 21^. Thus, Fyn might not directly interfere with TERT expression or telomerase activity. Rather, these data suggest that Fyn may interfere with endogenous telomere maintenance indirectly.

### Fyn phosphorylates Menin at tyrosine 603

After discovering that Fyn does not directly interfere with TERT expression or telomerase activity, we began to screen a candidate list of transcriptional modulators related to telomere maintenance (Supplementary Table 1)^22^. We included the transcriptional repressor Menin that is known to bind to telomeres^15, 22^. We performed Fyn immunoprecipitation with WT mESCs lysates using an anti-Fyn antibody and blotted against the candidate proteins. Of the candidate proteins, only the scaffold protein Menin showed significant interaction with Fyn (Supplementary Fig. 8). We later found that Menin is involved in telomere maintenance through interaction with TR and/or telomeres (see below). Endogenous Menin was detected in anti-Fyn-precipitated protein complexes in both E14 cells (Fig. 2A) and WT mESCs isolated from mice (Fig. 2B). We performed the previously described *in vitro* kinase assay with full-length recombinant human Menin purified from wheat germ expression system and found that Fyn does indeed phosphorylate Menin (Fig. 2C). Tandem mass spectrometry analysis identified Menin Y603 as the site of Fyn phosphorylation (Supplementary Fig. 9).

**Fig. 2.**
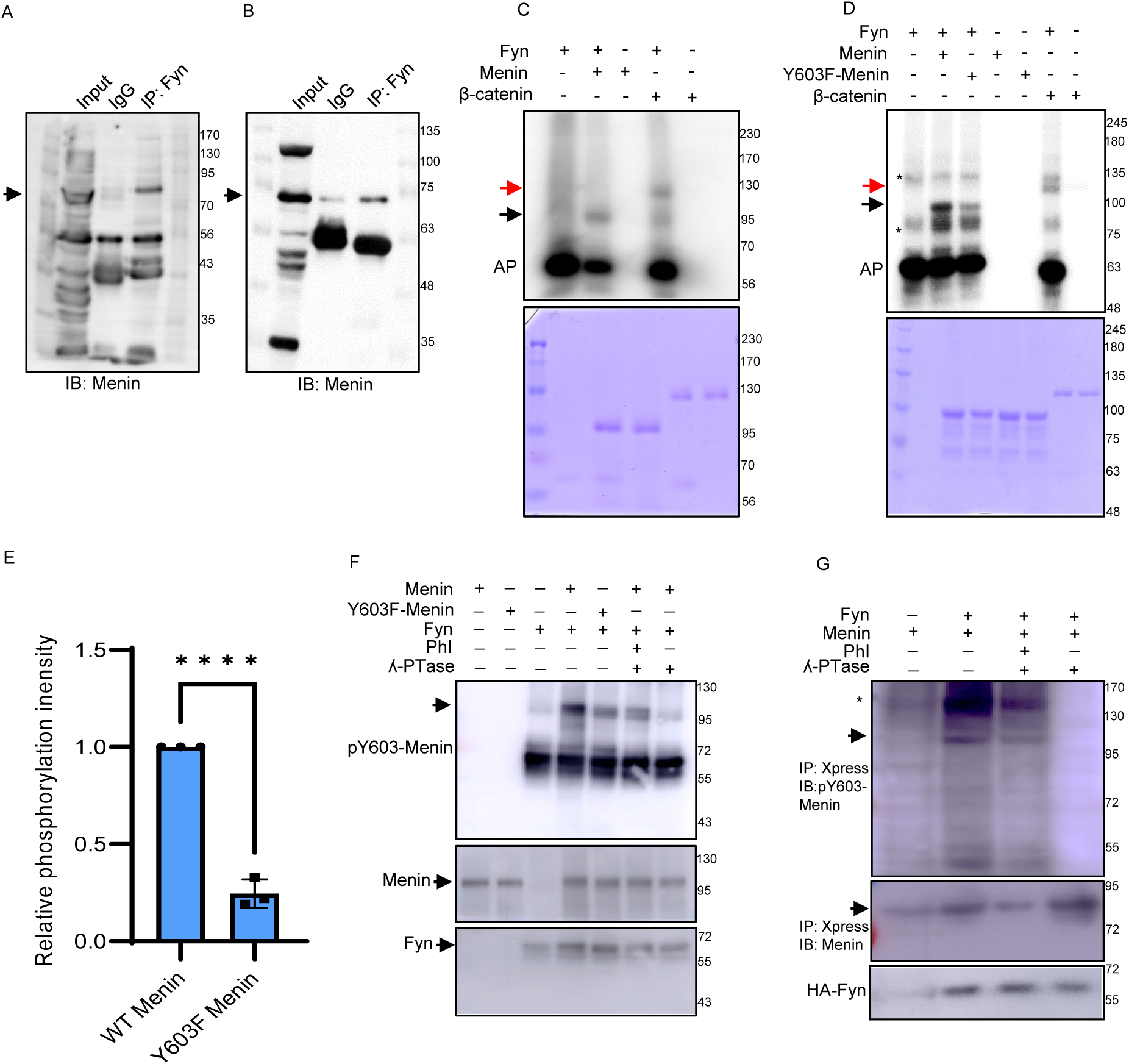
Fyn phosphorylates Menin at tyrosine 603. Immunoprecipitation using an anti-Fyn antibody followed by Western blotting using an anti-Menin antibody showing the interaction of endogenous Fyn and Menin in E14TG2a ES cells (A) and primary mESCs (B). Molecular weight markers are shown at the right. (C) Active Fyn and full-length Menin were subjected to an *in vitro* kinase assay with [γ-^32^P]ATP. Phosphorylated Menin was visualized by autoradiography indicated by an arrow. Full-length β-catenin was used as a positive control (red arrow). Menin and β-catenin were stained with Coomassie blue as loading controls (lower gel). AP= autophosphorylation. Molecular weight markers are shown at the right. (D and E) Fyn phosphorylates Menin at tyrosine 603. *In vitro* kinase assay (D) and quantification (E) of Fyn-mediated phosphorylation of full-length His-Menin and mutant His-Y603F-Menin in the presence of [γ-32P]ATP. β-catenin was used as a positive control of Fyn phosphorylation (red arrow). Phosphorylated Menin was detected by autoradiography, indicated by an arrow. Increased phosphorylation was detected in WT Menin compared to mutant Y603F Menin. The lower gel matches the key above to show WT Menin, Y603F Menin, and β-catenin stained with Coomassie blue. AP = autophosphorylation. * indicates nonspecific bands. Molecular weight markers are shown at the right. The phosphorylation band intensities of WT Menin and Y603F Menin were quantified using ImageJ. Data are presented as mean ± SD from a two-tailed t-test (**** = *p*LJ<LJ0.0001). A probability of *p* < 0.05 was considered statistically significant. (F) Phosphorylation of WT Menin by Fyn is reduced under phosphatase treatment. *In vitro* kinase assay of Fyn-mediated phosphorylation of WT His-Menin and mutant His-Y603F-Menin in the presence of 200 µM ATP and phosphatase. Phosphorylated Menin was detected by Western blot using a custom-synthesized antibody against (phospho-tyrosine 603-Menin) pY603-Menin. Fyn-mediated phosphorylation of Menin was blocked using Lambda phosphatase as indicated in the materials and methods section. The phosphatase effect was reversed using a phosphatase inhibitor (PhI). The arrow indicates phosphorylated Menin. * indicates nonspecific bands. The same membrane was stripped and reprobed using anti-Menin or anti-Fyn antibodies (lower blots). Molecular weight markers are shown at the right. (G) *In vivo* kinase assay of Xpress-Menin pulldown in HEK 293T cells. Xpress-Menin and HA-Fyn were over-expressed in HEK 293T cells. pY603-Menin was visualized by Western blotting using a custom-synthesized antibody against pY603-Menin. Lambda phosphatase was used to block Menin phosphorylation. The phosphatase effect was reversed using a PhI. (* indicates nonspecific bands). Pulldown of Xpress-tagged Menin and immunoblot against Xpress shows the presence of pull-down proteins in all lanes (middle blot). Pulldown of Xpress-tagged Menin and immunoblot against the HA-tagged antibody shows the presence of Fyn in Fyn-positive lanes (bottom blot).

We performed a radioactive *in vitro* kinase assay with active Fyn to compare WT and mutant Menin that cannot be phosphorylated at Y603 (Menin Y603F). These WT and mutant Menin proteins were purified using an *E. coli* expression system. The assay showed that phosphorylation of Menin Y603F is greatly decreased (Figs. 2D-E), confirming that Y603 serves as the site of Fyn phosphorylation.

We also performed a non-radioactive *in vitro* kinase assay with Fyn and WT or Y603F-Menin protein purified using an *E.coli* expression system. A custom-synthesized pY603-Menin antibody showed that phosphorylation of Menin Y603F is greatly decreased compared to WT Menin (Fig. 2F). To validate specificity of phospho-specific antibody, lambda phosphatase was used to eliminate phospho-signal. pY603-Menin signal was nearly abolished in the lambda phosphatase treated sample, but the signal was recovered by phosphatase inhibitor treatment (Fig. 2F).

In addition, we evaluated whether Fyn phosphorylates Menin at Y603 endogenously (*in vivo* kinase assay). We transfected HEK 293T cells with Xpress-tagged Menin to compare the amount of pY603-Menin in Xpress-Menin immunoprecipitation samples with or without Fyn over-expression. When co-expressed with Fyn, Xpress-tagged Menin showed pY603 signal that was not present in the sample without Fyn (Fig. 2G). Consistent with our previous observation, the pY603 signal was abolished in the lambda phosphatase-treated sample and recovered in the phosphatase inhibitor-treated sample. Together, these data suggest that Y603 serves as the site of Fyn phosphorylation and the custom synthesized antibody against pY603-Menin is specific to the site.

### Menin does not affect TERT transcription in mESCs

A previous study described Menin as a cell-type specific negative regulator of TERT transcription, but overexpression of Menin did not significantly reduce *in vitro* telomerase activity^23^. Although we found that Fyn deletion or inhibition did not affect TERT expression or *in vitro* telomerase activity (Supplementary Figs. 7A-F), we evaluated whether Fyn’s phosphorylation of Menin is involved with TERT transcription in stem cells. We performed a chromatin-immunoprecipitation assay using anti-Menin antibody in Sh-Mock vs Sh-Fyn E14 stem cells to evaluate whether Fyn-mediated phosphorylation of Menin affects its binding at TERT promoter. qPCR of the TERT promoter showed no significant difference in Menin occupancy between sh-Mock vs sh-Fyn E14 nuclear lysates (Supplementary Fig. 10A). These data suggest that Fyn-mediated phosphorylation of Menin does not regulate TERT transcription in stem cells. We also tested whether Menin binds to TERT. Immunoprecipitation results showed that Menin appears not to be a binding partner with TERT (Supplementary Fig. 10B).

### Fyn promotes Menin localization to telomeres

A previous study found that Menin localizes to telomeres in mouse spermatocytes during meiosis^15^. We proceeded to investigate whether Menin localizes to telomeres in stem cells by performing IF-FISH in unsynchronized mESCs with anti-Menin antibody and telomere FISH probe. We found that Menin does indeed co-localize with telomeres (Fig. 3A). Strikingly, Fyn^−/−^ mESCs showed a significant decrease in telomere/Menin co-localization compared to WT mESCs. These IF-FISH results suggest that Fyn-mediated phosphorylation of Menin promotes Menin localization to telomeres. It is not clear from these results whether Menin directly interferes with telomerase recruitment or activity at the site of telomere elongation.

**Fig. 3.**
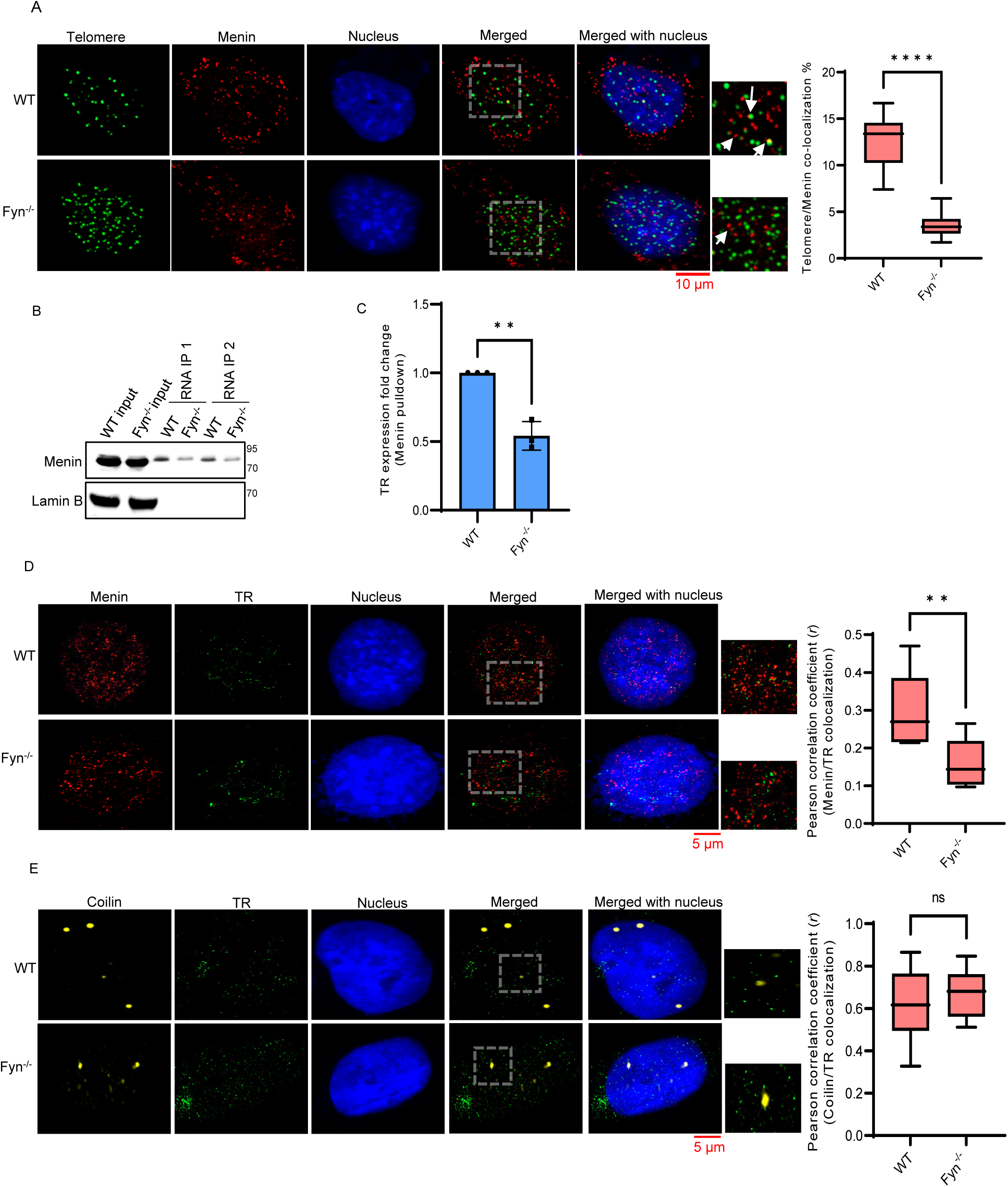
Fyn promotes Menin co-localization to telomeres and association with telomerase RNA *in vitro*. (A) IF-FISH analysis and co-localization analysis (B) of unsynchronized interphase cells in WT and Fyn^−/−^ mESCs shows Menin and telomere co-localization (scale 10 µm). Telomere/Menin co-localization was calculated manually. Both, totally merged and partially merged Telomere/Menin signals were considered co-localization counts. Data are presented as means of co-localization counts ± SD in 20 nuclei counts. Unpaired student’s t-test was used for statistical analysis (**** = *p*LJ<LJ0.0001). (B) Western blotting of *in vitro* transcribed Telomerase RNA (TR) pulldown assays using nuclear lysates from WT Fyn and Fyn^−/−^ mESCs shows Menin interaction with TR. Molecular weight markers are shown at the right. (C) RNP pulldown assay with anti-Menin antibody followed by qRT-PCR for TR in WT and Fyn^−/−^ mESCs shows increased TR amplification in WT vs Fyn^−/−^ mESCs. Unpaired student’s t-test was used for statistical analysis (** = *p*LJ<LJ0.01). (D) Representative IF-FISH images and quantification of Pearson correlation coefficient (*r*) of colocalization using an ImageJ plugin ‘Coloc 2’ shows significantly higher Menin (IF) and TR (FISH) binding in WT cells compared to Fyn^−/−^ cells. Data are presented as means ± SD from a two-tailed t-test (** = *p*LJ<LJ0.01). The scale bar is 5µm. (E) Representative of IF-FISH images and quantification of Pearson correlation coefficient (*r*) of colocalization using an ImageJ plugin ‘Coloc 2’ shows no significant difference in Coilin (IF) and TR (FISH) binding in WT cells compared to Fyn^−/−^ cells. The scale bar is 5 µm. Data are presented as means ± SD from a two-tailed t-test (ns, non-significant). A probability of *p* < 0.05 was considered statistically significant.

### Menin is associated with TR *in vitro* and its association with TR is modulated by Fyn

We next investigated whether Menin interacts with the telomerase RNA component (TR) using an *in vitro* RNA pulldown assay with *in vitro* transcribed biotinylated TR and nuclear lysates from mESCs. Western blotting from two independent experiments revealed that Menin was present in the TR pulldown complex. Interestingly, Menin showed greater association with TR in WT mESCs vs Fyn^−/−^ mESCs (Fig. 3B). We also performed Menin immunoprecipitation to compare TR expression in WT mESCs vs Fyn^−/−^ mESCs. TR expression was significantly decreased in the Menin immunoprecipitation sample from Fyn^−/−^ mESCs (Fig. 3C). These results show that Menin is a TR-associated protein and suggest that Fyn increases Menin’s association with TR. We performed IF-FISH with anti-Menin antibody and TR FISH probes in unsynchronized cells. The Pearson coefficient (*r*) of co-localization for TR and Menin was significantly higher in WT cells compared to Fyn^−/−^ cells (Fig. 3D), confirming that Fyn increases Menin’s association with TR. We also evaluated co-localization of TR and Coilin, a cajal body marker, in WT and Fyn^−/−^ mESCs (Fig. 3E). Although cajal bodies are not essential for telomerase biogenesis or activity^24, 25^, they are involved in telomerase trafficking and TR accumulates at cajal bodies in fixed cells^26–28^.

Interestingly, there was no difference in TR/Coilin co-localization between WT and Fyn^−/−^ mESCs (Fig. 3E). These results show that Fyn’s phosphorylation of Menin does not interfere with TR accumulation in cajal bodies and, taken together with TRAP assay results (Supplementary Figs. 7C-F), suggest that phospho-Menin does not interfere with telomerase assembly.

### Phosphorylation of Menin at Y603 promotes SUMO1 modification

Considering that many studies have demonstrated telomerase function and its subunits are modified by small ubiquitin-like modifier (SUMO) proteins^29–33^, we were curious to see if Menin and phospho-Menin are modified by SUMO proteins, which is known to affect stability and localization^34^. Interestingly, a phosphorylation-dependent SUMO modification consensus motif has previously been reported^35^. Another report showed SUMO modification of Menin at K591 and suggested that Menin contains multiple SUMOylation sites^36^. To further study Menin SUMOylation, we used GPS-SUMO (http://sumosp.biocuckoo.org/) to identify additional SUMOylation sites^37^. We selected lysine residues with the lowest p-values, 609 (K609) as possible SUMO-modification site (Supplementary Fig. 11A) and proceeded to investigate whether Fyn’s phosphorylation of Menin at Y603 regulates its SUMOylation. We performed immunoprecipitation with anti-Menin antibody and blotted against SUMO1 and SUMO2/3/4. Results showed that endogenous Menin interacted with SUMO1 in Sh-Mock cells, but the interaction was greatly reduced in Sh-Fyn cells (Fig. 4A-B), suggesting that Fyn-mediated phosphorylation of Menin is important for SUMO1 interaction with Menin. We did not observe an interaction between endogenous Menin and endogenous SUMO2/3/4 (Supplementary Fig. 11B). We also used an ELISA SUMOylation assay that specifically detects SUMO1 to measure Menin SUMO1 modification in nuclear lysates from mESCs. We observed a reduced SUMOylation intensity in Fyn-knockout cells vs. WT cells. Moreover, SUMOylation intensity was also reduced in Sh-Fyn compared to Sh-Mock mESCs (Supplementary Fig. 11C).

**Fig. 4.**
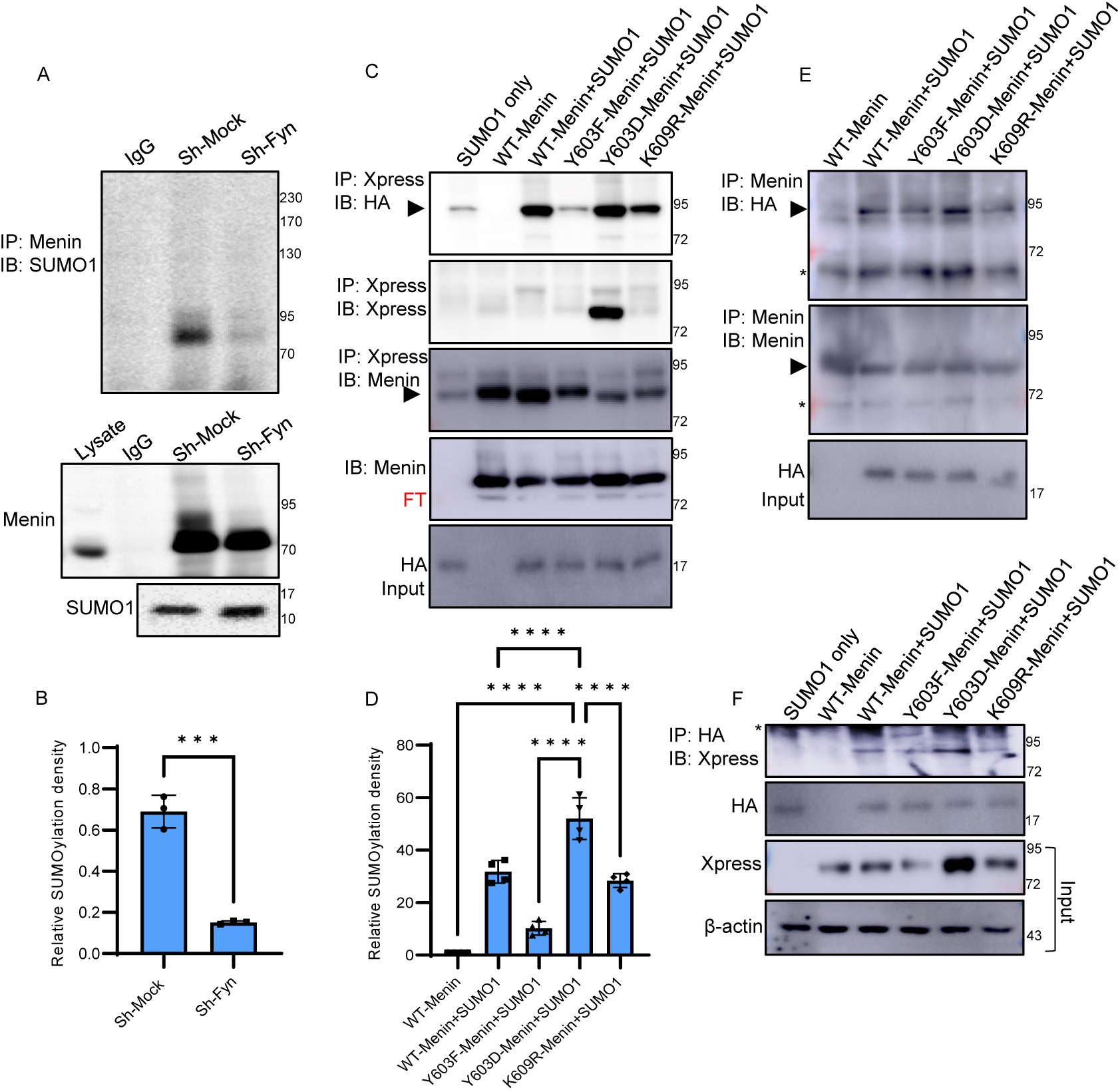
Phosphorylation of Menin at Y603 promotes SUMO1 modification at K609. (A) Endogenous Menin interacts with SUMO1 in the presence of Fyn. Immunoprecipitation of Menin and Western blotting of SUMO1 in sh-Mock and sh-Fyn E14 cells (upper). Endogenous expression of Menin in immunoprecipitated samples (middle) and SUMO1 as input (lower). Molecular weight markers are shown at the right. (B) Endogenous SUMOylation intensity was quantified using ImageJ. Menin densitometry was used to normalize SUMOylation intensity. Data are presented as means ± SD from a two-tailed t-test (*** = *p*LJ<LJ0.001). (C) Mutations at tyrosine 603 and lysine 609 impair SUMOylation of Menin as shown by immunoprecipitation of overexpressed WT or mutant Xpress-tagged Menin and immunoblotting of HA-tagged SUMO1 in HEK 293T cells. Phosphodeficient Menin mutant (Y603F) and a SUMO-deficient Menin mutant (K609R) show reduced SUMOylation, whereas the phosphomimetic mutant Y603D Menin mutant show increased SUMOylation. Molecular weight markers are shown at the right. Black arrow, SUMO1-Menin, red arrow, Menin; FT, flowthrough. (D) Quantification of SUMOylation intensity in WT, Y603F, Y603D, K609R Menin using ImageJ from Fig. 5C. Menin densitometry was used to normalize SUMOylation intensity. Data are presented as means ± SD from a One-way ANOVA followed by Tukey’s multiple comparison tests (**** = *p*LJ<LJ0.0001). (E) Mutations at tyrosine 603 and lysine 609 impair SUMOylation of Menin as shown by immunoprecipitation of overexpressed WT or mutant Menin and immunoblotting of HA-tagged SUMO1 in HEK 293T cells. Phosphodeficient Menin mutant (Y603F) and a SUMO-deficient Menin mutant (K609R) showed reduced SUMOylation, whereas the phosphomimetic mutant Y603D Menin mutant showed increased SUMOylation. (F) Phosphodeficient Menin mutant (Y603F) and a SUMO-deficient Menin mutant (K609R) showed reduced SUMOylation, whereas the phosphomimetic mutant Y603D Menin mutant showed increased SUMOylation as shown by pulldown of HA-tagged SUMO1 and immunoblotting of Xpress-tagged Menin in HEK 293T cells. A probability of *p* < 0.05 was considered statistically significant.

To gain insight into the mechanism of Fyn-mediated Menin SUMOylation, we transfected HEK 293T cells with Xpress-tagged pcDNA-WT-Menin (WT*-*Menin), pcDNA-Y603F-Menin (Y603F-Menin), and pcDNA-Y603D-Menin (Y603D-Menin) with or without pcDNA-HA-SUMO1 (SUMO1). Menin Y603D served as a phosphomimetic mutant. We performed Xpress immunoprecipitation followed by western blot with anti-HA antibody. No SUMO1-Menin band was observed in cells transfected with WT-Menin only (Fig. 4C, upper blot). SUMO1-Menin was observed in both WT-Menin+SUMO1 and Y603D-Menin+SUMO1-transfected cells, but SUMO1-Menin was reduced significantly in Y603F-Menin+SUMO1*-*transfected cells (Fig. 4C-D). The significantly reduced SUMO1-Menin band in Y603F-Menin+SUMO1-transfected cells suggests that SUMOylation modification of Menin is phosphorylation-dependent (Fig. 4C-D). Interestingly, K609R-Menin+SUMO1-transfected cells show greater SUMO1-Menin signal than Y603F-Menin+SUMO1-transfected cells, indicating that Y603 phosphorylation may affect SUMO1 modification at additional residues. Surprisingly, anti-Xpress reblotting of Xpress IP membrane with Xpress antibody shows a very strong Xpress-Menin band in Y603D-Menin*+*SUMO1 transfected cells compared to WT*-*Menin*+*SUMO1 and other mutants (Fig. 4C). However, reblotting of the same membrane with Menin antibody did not show any difference between Y603D-Menin*+*SUMO1, Y603F-Menin*+*SUMO1 and K609R-Menin*+*SUMO1 (Fig. 4C). These results suggested that the C-terminal region of Xpress-Menin, which contains the Xpress tag and mutation sites, may be subject to degradation and that phosphorylation at 603 prevents C-terminal degradation. Later we explore this mechanism in more detail (see Fig. 5 and Supplementary Fig. 12).

**Fig. 5.**
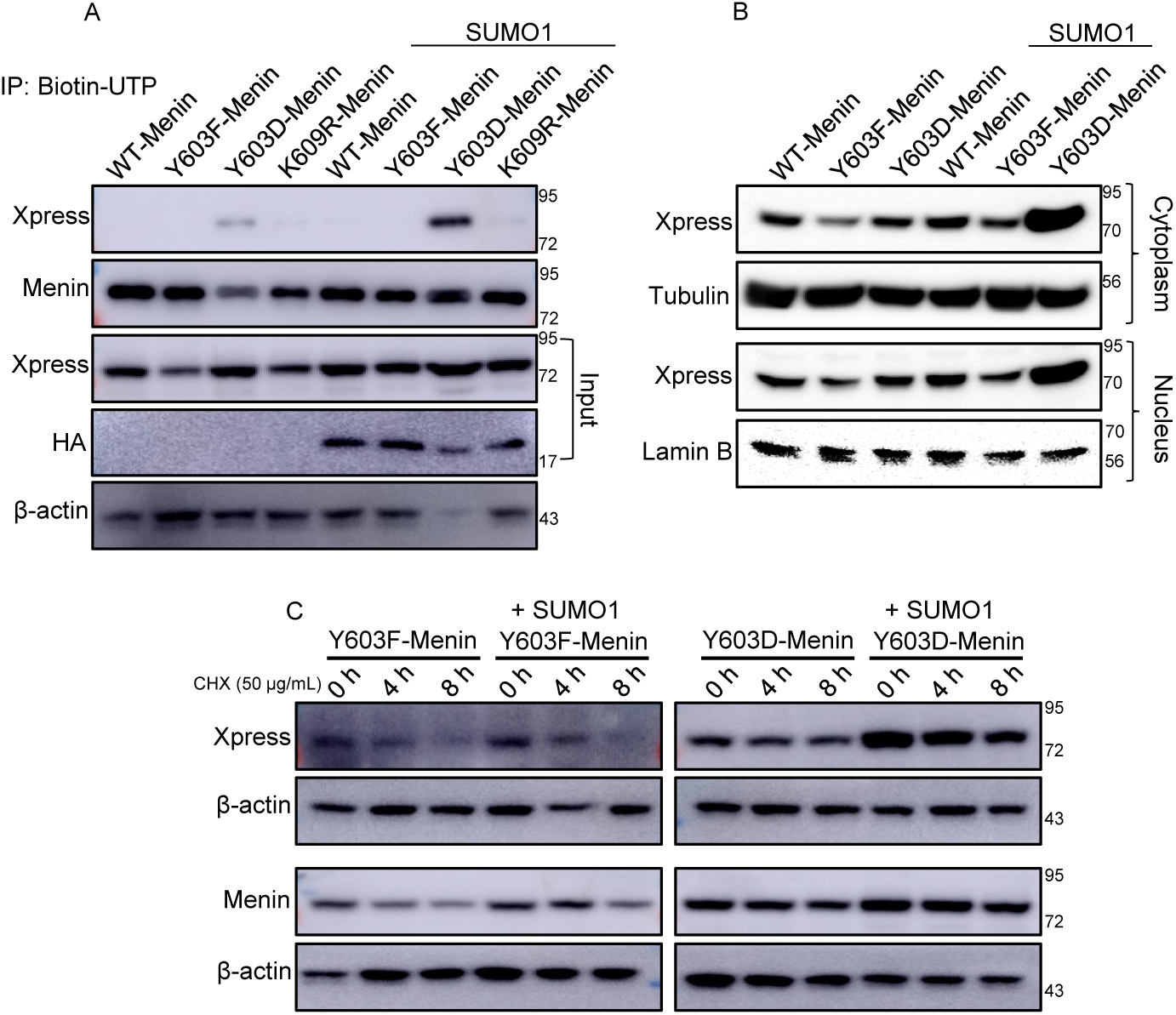
Menin Phosphorylation and SUMO1 modification promote Menin association with telomerase RNA by increasing protein stability. (A) Phosphorylation and SUMO1 modification of Menin promote Menin-TR association. RNA pulldown assay using *in vitro* transcribed TR and overexpressed WT or mutant (Y603F, Y603D, K609R) Menin with *or* without SUMO1. Menin was analyzed with an Xpress antibody that binds the C-terminal Xpress Tag of the protein or a Menin antibody that binds the central region of Menin. The Western blot using Xpress antibody indicates that Y603D Menin with SUMO1 expression is greatly increased in TR pulldown complex compared to WT, Y603F and K609R Menin with or without SUMO1 expression. Y603D-Menin shows less expression in TR pulldown binding than Y603D-Menin+SUMO1. No significant changes were detected with Menin antibody. We intentionally loaded less amount of input in the Y603D-Menin+SUMO1 lane to visualize bands in all lanes. (B) Western blotting of cytoplasmic and nuclear protein extracts shows that phosphomimetic Y603D-MEN1 and SUMO1 promote Menin stability. HEK 293T cells were transfected with Xpress-tagged *MEN1* (WT or Y603F, Y603D) with or without HA-tagged SUMO1. (C) Cycloheximide chase assay showing significantly higher protein degradation in Y603F-Menin with or without SUMO1 compared to Y603D-Menin with or without SUMO1. Y603F and Y603D *MEN1*transfected HEK 293T cells co-transfected with *SUMO1* were treated with 50µg/mL cycloheximide for indicated time points.

In a similar experimental model, we performed Menin pulldown using anti-Menin antibody with lysates from WT+SUMO1 or mutant Menin+SUMO1 co-transfected HEK 293T cells. Fig. 4E (upper blot) shows more HA-SUMO1 band in Y603D-Menin+SUMO1-transfected cells compared to Y603F-Menin+SUMO1 and K609R*-*Menin+SUMO1 transfected cells. In an additional experimental model, we pulled down HA-SUMO1 with HA antibody using the same cell lysates that were used in previous co-transfection experiments. The anti-Xpress immunoblot in Fig. 4F shows increased Xpress-Menin in HA pulldown from Y603D-Menin+SUMO1-transfected cells compared to HA pulldowns from WT+SUMO1, Y603F-Menin+SUMO1, and K609R-Menin+SUMO1-transfected cells. Taken together, these immunoprecipitation results suggest that phosphorylation of Menin at Y603 promotes SUMO1 modification.

### Phosphorylation-dependent SUMO1 modification of Menin promotes Menin’s association with TR by increasing Menin stability

In order to determine if SUMO1 modification of Menin affects Menin/TR association, we performed TR pulldown assays with HEK 293T lysates co-overexpressing WT or mutant Menin with or without SUMO1. Interestingly, we detected significantly higher Xpress-Menin in TR pulldown complexes of Y603D-Menin+SUMO1-transfected cells (Fig. 5A). The Y603D-Menin without SUMO1 sample showed Xpress-Menin/TR association, but to a lesser extent than Y603D-Menin+SUMO1 sample (Fig. 5A). Y603F-Menin and K609R-Menin samples show negligible or no binding with TR irrespective of SUMO1 transfection (Fig. 5A). These results suggest that phosphorylation and SUMOylation of Menin at the C-terminal domain promote its association with TR. Conversely, reblotting with an anti-Menin antibody that recognizes the center region of Menin, showed no change between WT and mutant Menin with or without SUMO1 (Fig. 5A). Taken together, the seemingly contradictory results of anti-Xpress and anti-Menin blots suggest that there are at least two distinct domains of Menin associated with TR. To further investigate the strong binding affinity between TR and Y603D-Menin*+*SUMO1 lysates, we performed a modified TR-pulldown experiment, where 50-fold less Y603D-Menin+SUMO1 input was used compared to Y603F-Menin+SUMO1 and K609R-Menin+SUMO1 input. The input lanes show that Xpress-Menin expression is near equal between the samples, while β-actin expression shows a 50-fold difference (Supplementary Fig. 12A). In TR-IP lanes, even 50-fold less input from Y603D-Menin+SUMO1 sample shows increased TR binding affinity compared to Y603F-Menin+SUMO1 and K609R-Menin+SUMO1.

We assessed Menin subcellular expression and localization comparing WT, Y603F, and Y603D Menin with or without SUMO1 in cytoplasmic and nuclear extracts. The ratio of cytoplasmic to nuclear Menin expression did not change between WT and mutant Menin (Fig. 5B). These results indicate that phosphorylation-dependent SUMOylation of Menin promotes its stability, but does not affect subcellular localization.

To further explore how the C-terminal phosphorylation and SUMO1 modification affect Menin stability, we performed a Cyclohexamide (CHX) Chase Assay in Y603F-Menin and Y603D-Menin with or without SUMO1. The Xpress-Menin shows a time dependent degradation in both Y603F-Menin and Y603D-Menin, but the protein turnover was much slower in Y603D-Menin, likely due to increased stability at the C-terminal region (Fig. 5C, Supplementary Fig. 12B). The anti-Menin antibody that recognizes the central region shows that stability of Y603D-Menin was slightly higher in the presence of SUMO1 compared to Y603F-Menin+SUMO1 (Fig. 5C, Supplementary Fig. 12B). The stability of both Y603F-Menin and Y603D-Menin was higher in the presence of SUMO1, indicating that SUMO1-modification increases Menin stability. In order to directly compare Menin stability between Y603D and Y603F, we performed western blotting with CHX assay lysates on the same membrane. A time dependent exposure of western blot membrane showed no Xpress-Menin in Y603F-Menin lysates after 1 minute, while Xpress Menin showed significant expression in Y603D-Menin lysates. Xpress-Menin was only visible in the Y603F-Menin lysates after a 10-minute exposure (Supplementary Fig. 12B). Detection of the central region of Menin shows Menin expression at 1 min or 10 min of exposure in both Y603F-Menin and Y603D-Menin lysates (Supplementary Fig. 12C). However, Menin shows much greater stability in Y603D-Menin lysates compared to Y603F-Menin lysates. Altogether, these results suggest that phosphorylation and SUMOylation at the C-terminal region of Menin increase its stability and/or promote resistance to C-terminal degradation, ultimately promoting Menin’s association with TR.

### Overexpressed, SUMO1-modified Menin inhibits TR association with Dyskerin and GAR1

We performed tandem mass spectrometric analysis of TR pull-down complex from Y603F-Menin+SUMO1 and Y603D-Menin+SUMO1 co-transfected HEK 293T nuclear lysates. The results indicated that 54 human proteins were associated with TR in Y603F-Menin-transfected cells. The number of proteins was reduced dramatically in Y603D-Menin+SUMO1 co-transfected cells (Fig. 6A and Supplementary Table 2). Importantly, Y603D-Menin transfection inhibited TR association with telomerase subunits Dyskerin and GAR1 (Fig. 6B and Supplementary Table 2).

**Fig. 6.**
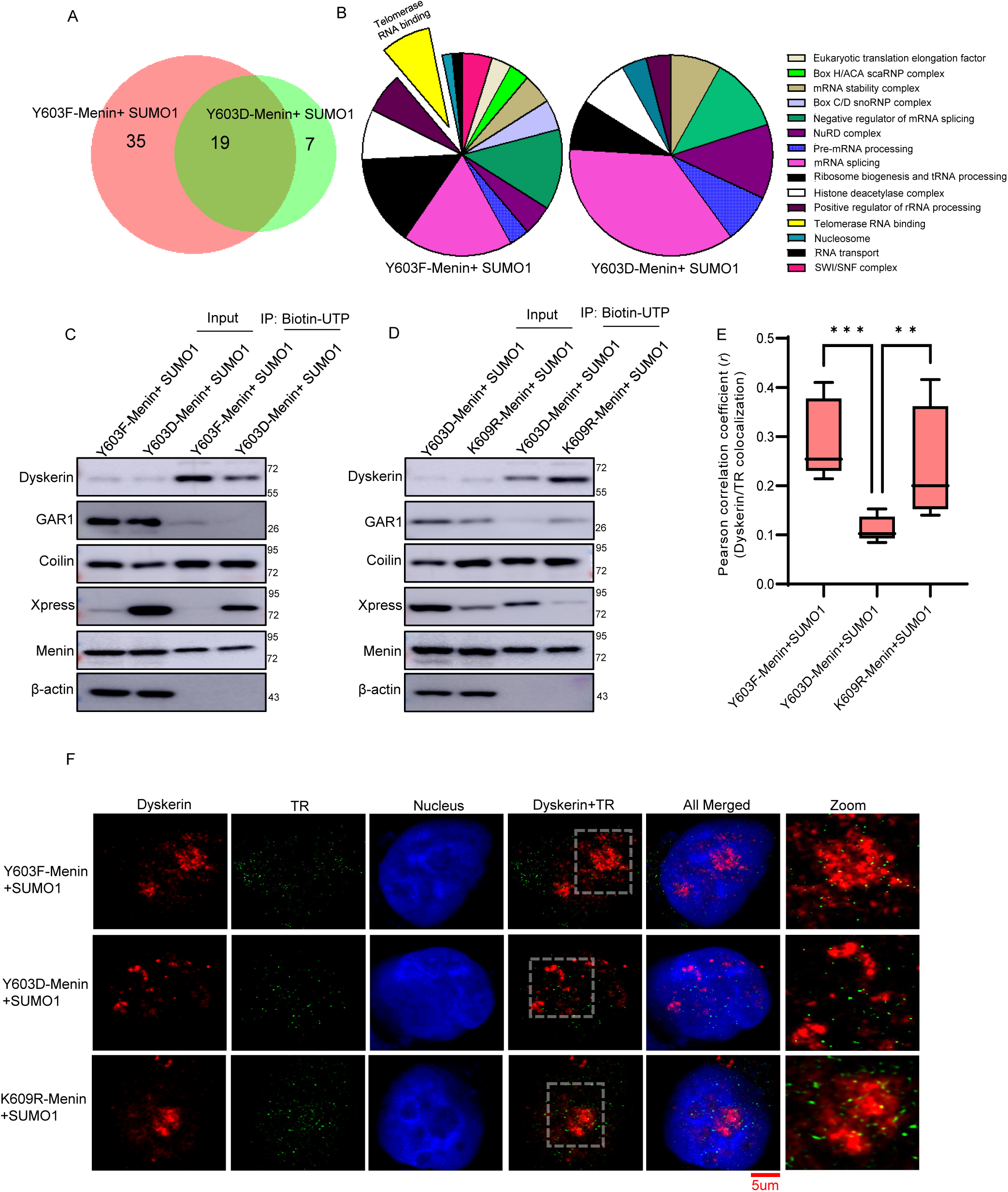
SUMO1-modified Menin prevents recruitment of Dyskerin and GAR1 to TR. (A) A Venn diagram determined by mass spectrometry analysis displays differential proteins recruited to TR after Y603F-*MEN1*+*SUMO1* or Y603D-*MEN1*+*SUMO1* transfection in 293T cells. (B) A Pie-diagram shows classes of TR binding proteins in Y603F-*MEN1*+*SUMO1*– and Y603D-*MEN1*+*SUMO1*-transfected 293T cells. (C) Western blotting of *in vitro* transcribed TR pulldown assay from whole cell lysates of HEK 293T cells with over-expressed mutant (Y603F, Y603D) Menin and SUMO1 shows decreased Dyskerin and GAR1 binding in Y603D-*MEN1*+*SUMO1* transfected cells. Coilin binding shows no difference between Y603F vs Y603D. Xpress-tagged Menin shows increased binding in Y603D-Menin compared to Y603F-Menin although Menin expression shows no difference. Molecular weight markers are shown at the right. (D) Western blotting of *in vitro* transcribed TR pulldown assay from whole cell lysates of HEK 293T cells with over-expressed mutant (Y603D, K609R) Menin and SUMO1 shows increased Dyskerin and GAR1 binding in K609R-*MEN1*+SUMO1 transfected cells. Coilin binding shows no difference between Y603D vs K609R. Xpress-tagged Menin shows increased binding in Y603D-Menin compared to K609R-Menin although Menin expression shows no difference. Molecular weight markers are shown at the right. (E, F) Quantification of the Pearson correlation coefficient (*r*) of colocalization using an ImageJ plugin ‘Coloc 2’ between representative IF-FISH images shows significantly less Dyskerin and TR binding in Y603D-*MEN1*+SUMO1 transfected 293T cells compared to mutant Y603F-*MEN1*+*SUMO1* and K609R-*MEN1+*SUMO1 transfected cells. The scale bar is 5 µm. Data are presented as means ± SD from a one-way ANOVA followed by Tukey’s multiple comparison tests (** = *p*LJ<LJ0.01, *** = *p*LJ<LJ0.001). A probability of *p* < 0.05 was considered statistically significant.

We corroborated mass spectrometry results by performing TR-IP followed by Western blot using Y603F-Menin+SUMO1 and Y603D-Menin+SUMO1 co-transfected HEK 293T lysates. Whole cell lysates showed that Dyskerin and GAR1 levels were not reduced, suggesting that Menin does not affect Dyskerin or GAR1 expression (Fig. 6C). However, Dyskerin and GAR1 binding were reduced in TR-IP of Y603D-Menin+SUMO1 transfected cells, confirming the TR mass spectrometry results (Fig. 6C). Coilin/TR binding was not changed between these two groups (Fig. 6C). We also compared Dyskerin and GAR1 expression in TR-IP of whole cell lysates of 293T cells transfected with Y603D-Menin+SUMO1 or K609R-Menin+SUMO1. We found that cells transfected with SUMO-deficient Menin (K609R), showed greater Dyskerin and GAR1 binding in TR-IP (Fig. 6D). Xpress-tagged Menin binding was also reduced in the K609R sample, further supporting that SUMO1 modification of Menin at K609 promotes its association with TR, Menin blot did not show any changes further confirming our previous results (Fig. 6D).

We also performed IF-FISH experiments with DKC1 antibody and TR FISH probe to evaluate co-localization of Dyskerin and TR after Menin and SUMO1 overexpression. IF-FISH analysis showed significantly more Dyskerin and TR co-localization in Y603F-Menin+SUMO1 and K609R-Menin+SUMO1 compared to Y603D-Menin+SUMO1 transfected 293T cells (Fig. 6E, F). Transfection of Menin mutants did not affect the binding of Coilin to TR (Supplementary Fig. 13). Collectively, these results suggest that association of SUMO1-modified Menin with TR restricts Dyskerin’s association with TR.

### Menin knockdown promotes telomerase localization to telomeres and slows telomere erosion

We performed Menin knockdown in HEK 293T cells to evaluate whether Menin expression affects telomerase assembly, telomerase localization to telomeres, and telomere length. Western blotting analysis in Sh-Mock vs Sh-Menin 293T cells confirmed efficient Menin knockdown (Fig. 7A). IF staining of Dyskerin and Coilin alongside FISH with TR probes showed no significant difference in either Coilin-Dyskerin co-localization or Coilin-TR co-localization (Fig. 7B-D) between Sh-Mock cells and Sh-Menin cells. These results further support that Menin does not interfere with telomerase assembly in Cajal bodies. Analysis of Dyskerin IF and telomere FISH showed that co-localization of Dyskerin and telomeres was increased in Sh-Menin 293T cells compared to Sh-Mock cells (Fig. 7E-F). These results suggest that Menin inhibition significantly increased Dyskerin localization to telomeres. A significantly higher Dyskerin/TR co-localization was also noted in Sh-Menin cells than Sh-Mock (Fig. 7G). Telomere length analysis by qRT-PCR showed significantly higher telomere to *Alu* sequence ratio (T/S) in Sh-Menin cells compared to Sh-Mock cells at both passages 5 and 10 (Fig. 7H). These studies indicate that Menin knockdown slows telomere erosion in a manner similar to Fyn Knockdown.

**Fig. 7.**
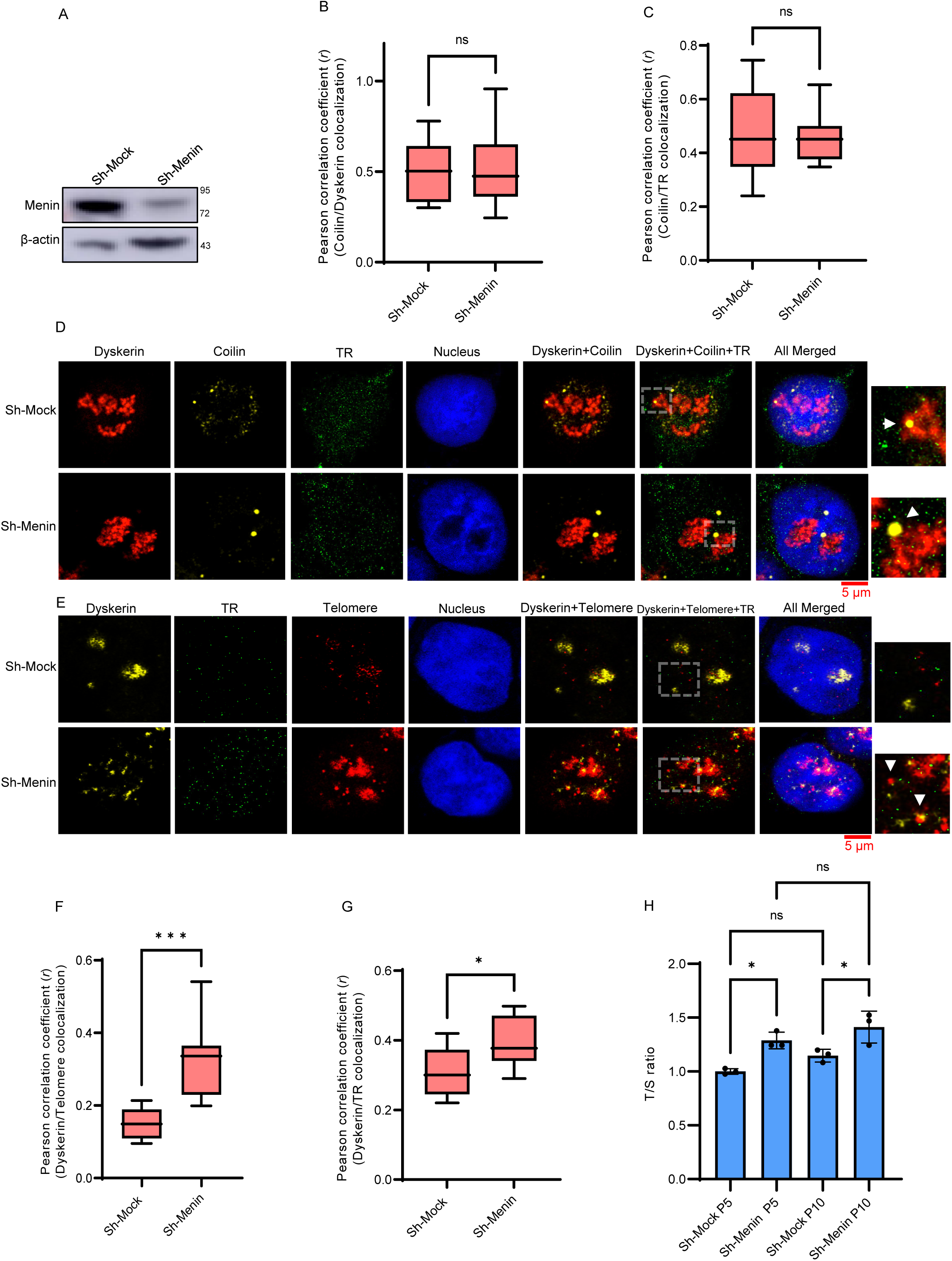
Menin knockdown improves Dyskerin localization to telomeres and telomere length without affecting Dyskerin accumulation in Cajal bodies. (A) Western blotting of Menin in sh-Mock and sh-Menin HEK 293T cells. β-actin served as a loading control. Molecular weight markers are shown at the right. Quantification of the Pearson correlation coefficient (*r*) for Coilin/Dyskerin (B) and Coilin/TR (C) co-localization analysis obtained by IF of Dyskerin, Coilin, and FISH of TR (D). IF quantitation data using ImageJ plugin ‘Coloc 2’ show no significant difference between Sh-Mock and Sh-Menin HEK 293T cells. Data are presented as means ± SD from a two-tailed t-test (ns, non-significant). Arrows indicate colocalization between DKC and Coilin. The scale bar is 5µm. Region of interest (ROI) was used to analyze Coilin/Dyskerin co-localization. (E, F, G) Dyskerin IF and TR/Telomere FISH and quantification of Pearson correlation coefficient (*r*) of colocalization using an ImageJ plugin ‘Coloc 2’ shows significantly more Dyskerin/telomere co-localization (F) and Dyskerin1/TR co-localization (G) in sh-Menin cells compared to sh-Mock HEK 293Tcells. Data are presented as means ± SD from an unpaired student’s t-test (* = *p*LJ<LJ0.05, *** = *p*LJ<LJ0.0001). Arrows indicate colocalization between DKC and Telomeres. The scale bar is 5µm. ROI was used to analyze Dyskerin/telomere co-localization. Clump telomere signals were avoided for quantifications. (H) Telomere length in sh-Menin 293T cells and sh-Mock cells as determined by quantitative PCR at passages 5 and 10. T/S ratios were calculated relative to the single-copy gene *Alu*. Data are presented as means ± SD from a one-way ANOVA followed by Tukey’s multiple comparison tests (* = *p*LJ<LJ0.05). A probability of *p* < 0.05 was considered statistically significant.

### Fyn knockdown in Human DC iPSCs slows telomere shortening

We next investigated Fyn inhibition in iPSCs from a DC patient that exhibit accelerated telomere erosion and telomerase deficiency due to a mutation in DKC1. We first used DC fibroblasts to generate DC iPSCs. iPSC generation was confirmed by the expression of SSEA-4, Tra-1-60, KLF4 and Nanong (Supplementary Fig. 14A-B). Next, we created DC iPSCs with Fyn KO using CRISPR-Cas9. A mock CRISPR-Cas9 plasmid was used as negative control. CRISPR-Cas9 transduction generated a population of iPSCs with significantly decreased Fyn expression, although expression was not completely abolished in the whole iPSC population (Fig. 8A). Therefore, we used the term Fyn knockdown (Fyn^KD^) to describe these iPSCs instead of Fyn knockout (Fyn^KO^). CRISPR-Cas9-mediated Fyn^KD^ reduced both Fyn and pY603-Menin expression in DC iPSCs (Fig. 8A-C). iPSCs showed no significant difference in telomere length after 3 passages (Fig. 8D-E). However, at passage 12, Fyn^KD^ iPSCs showed significantly longer telomeres than mock iPSCs as determined by Q-FISH (Fig. 8D, F). Consistent with these results, Flow-FISH and qRT-PCR results showed that Fyn^KD^ DC iPSCs exhibited significantly higher telomere signals compared to control DC iPSC after 12 passages (Fig. 8G and H, respectively).

**Fig. 8.**
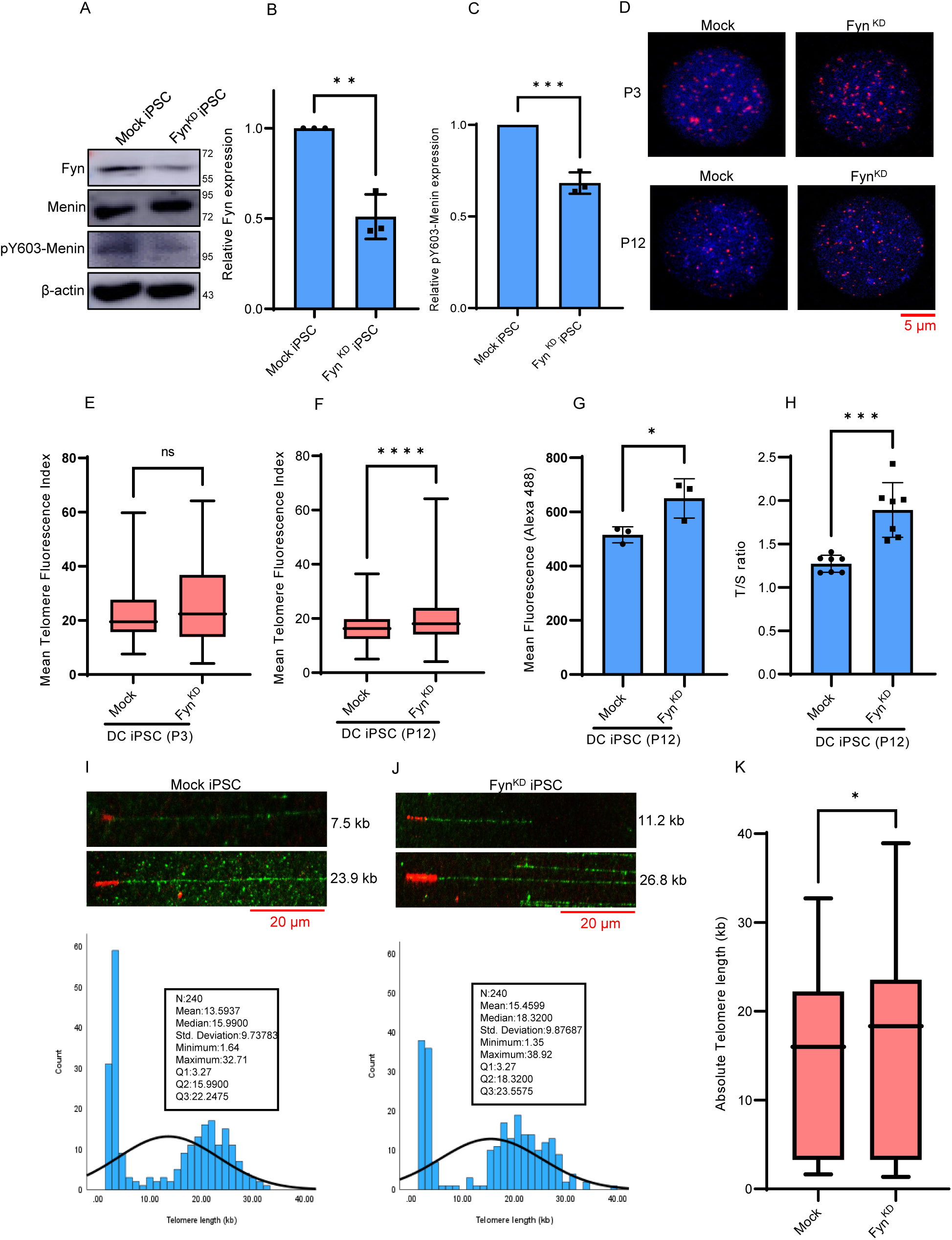
Fyn knockdown in Human DC iPSCs slows telomere erosion. (A) Western blotting showing *Fyn* gene knockdown efficiency in DC iPSCs using CRISPR-Cas9– and the expression of Menin and pY603-Menin. β-actin served as a loading control. Molecular weight markers are shown at the right. (B, C) Densitometry by ImageJ shows knockdown of Fyn (B) and reduced expression of pY603-Menin (C) in Fyn^KD^ DC iPSC vs mock DC-iPSC. Data are presented as means ± SD from a two-tailed t-test (** = *p*LJ<LJ0.01, *** = *p*LJ<LJ0.001). (D, E, and F) Representative images of interphase Q-FISH analysis (D) and quantification (E) reveal no significant differences in telomere length at passages 3 of Fyn^KD^ DC iPSCs vs mock DC iPSCs, but at passages 12 (F), Fyn^KD^ iPSCs showed higher Q-FISH signal vs mock iPSC (n = 50 nuclei). Data are presented as means ± SD from a two-tailed t-test (ns = not significant, **** = *p*LJ<LJ0.0001). (G) Flow-FISH analysis showed higher telomeric mean fluorescence in Fyn^KD^ DC iPSCs vs mock-DC-iPSCs. (H) qRT-PCR analysis shows a significantly higher T/S ratio in Fyn^KD^ DC iPSCs vs mock-DC-iPSCs. The human *Alu* gene was used for normalization. Data are presented as means ± SD from a two-tailed t-test (* = *p*LJ<LJ0.05, *** = *p*LJ<LJ0.001). (I, J) Representative images of telomere analysis using a Molecular Combing assay from genomic DNA fibers isolated from Mock iPSC vs Fyn^KD^ iPSC (passage 12). The scale bar is 20 µm. A detailed statistical analysis and frequency distribution are shown below each sample DNA fiber image. Q1, quartile 1; Q2, quartile 2 and Q3, quartile 3. (K) Absolute Telomere length in kb (n= 240 DNA fibers) between mock iPSC vs Fyn^KD^ iPSC. Data are presented as means ± SD from a two-tailed t-test (* = *p*LJ<LJ0.05). A probability of *p* < 0.05 was considered statistically significant.

Moreover, TMCA results also show that Fyn knockdown slows telomere erosion in DC iPSCs. DNA fibres from Fyn^KD^ iPSC show significantly longer telomeres than Mock iPSC (Fig. 8I-K). Detail statistical analysis from the telomere signals shows that mean and median telomere sizes were 15.4599 kb and 18.3200 kb, respectively, in Fyn^KD^ cells compared to Mock cells, which shows that mean and median telomere sizes were 13.5937 kb and 15.990 kb, respectively (Fig. 8I-K). A comparative statistical analysis shows significantly longer absolute telomere length in Fyn^KD^ cells compared to Mock cells (Fig. 8K).

## Discussion

This work uncovers a new mechanism of telomere maintenance in mammalian stem cells. We demonstrate that Fyn, an Src family kinase, is a negative regulator of mammalian telomere length. We first discovered that inhibition or deletion of Fyn stabilizes telomere length. However, we found that Fyn does not affect TERT expression and does not interfere with telomerase activity using *in vitro* assays. Rather, our results suggest Fyn negatively regulates telomere length by phosphorylating the scaffold protein Menin at Y603, promoting telomerase subunit mislocalization, and may compromise telomerase activity *in vivo*.

Phosphorylation of Menin at Y603 promotes phosphorylation-dependent SUMOylation at K609, which increases Menin’s C-terminal stability and association with TR. Importantly, phospho-site mutant Y603F and SUMO-site mutant K609R Menin showed decreased SUMOylation and stability, thereby reducing its association with TR. We found that SUMO1-modified Menin associates much more strongly with TR, decreasing TR’s association with telomerase subunits Dyskerin and GAR1.

Telomerase stability, assembly, and recruitment to telomeres are essential steps during telomere elongation^38^. Our results strongly suggest that knockdown of Fyn or Menin does not affect telomerase assembly, but rather, impairs telomerase holoenzyme function. One possible mechanism is that binding of Menin to TR promotes the dissociation of Dyskerin and GAR1 from telomerase RNPs en route to telomeres. An elegant study by Schmidt *et al* used live cell imaging to reveal that more than 85 % of telomerase RNPs are freely diffusing throughout the nucleus^28^. Our results point to the possibility that SUMO1-Menin compromises telomerase function by binding to TR in a portion of the telomerase RNP population, rather than the telomerase RNP accumulated in cajal bodies. Another possibility is that SUMO1-Menin compromises telomerase function by associating with TR at telomeres.

Defective telomere maintenance may result in a TBD such as DC^6, 39, 40^. We saw that CRISPR-Cas9-mediated Fyn inhibition in mutant DKC1 human DC iPSCs stabilized telomeres concurrent with a decline in pY603-menin expression. There are conflicting findings of telomere shortening in DC iPSCs. Agarwal *et. al.,* 2010 showed that DC iPSCs increased telomere length over time^41^. However, Batista et. al., Woo et al., and Fernandez et al observed telomere shortening in DC iPSCs over time. Consistent with the latter publications, we found telomere shortening in later passages of DC iPSCs accompanied by spontaneous differentiation^42–44^. It is tempting to speculate that endogenous Fyn exacerbates telomere shortening in these telomerase-defective human iPSCs by phosphorylating Menin and further compromising *in vivo* telomerase holoenzyme function. Indeed, our human DC iPSC data suggests that endogenous Fyn negatively regulates telomere maintenance even in the presence of defective telomerase.

In conclusion, our results demonstrate that Fyn-mediated phosphorylation and subsequent SUMO1-modification of Menin disrupt telomere maintenance in proliferating stem cells. We found that Fyn inhibition promotes telomere maintenance, even in telomerase-defective stem cells. Fyn inhibitors may be a viable treatment option for DC or other TBDs, although the development of specific inhibitors is needed.

## Materials and Methods

The antibodies, primers, kits, and chemicals used in this study are in Supplementary Tables 3-8. Plasmid Men1-pDNR-Dual (Cat. no: HsCD00000838) was purchased from DNASU. pcDNA3-HA-Sumo1 was a gift from Junying Yuan (Addgene plasmid # 21154)45.

### Fyn ^−/−^ animal model

Fyn^−/−^ mice (B6.129S7-Fyntm1Sor/J) were purchased from The Jackson laboratory^46^ and bred, propagated, and maintained at The Hormel Institute, University of Minnesota. Mice were maintained following the guidelines established by the University of Minnesota Institutional Animal Care and Use Committee. Fyn genotyping was performed by PCR using a protocol as recommended by The Jackson Laboratory. Representative genotyping characterization is shown in Supplementary Fig. 9A. For this study, age-matched wildtype (WT) and Fyn^−/−^ mice (n = 12 males + 12 females) were maintained for 97 weeks. Some mice needed to be euthanized based on the veterinary advice. This study was completed with 13 WT and Fyn ^−/−^ mice (6 males + 7 females).

### Generation of mouse embryonic stem cells (mESCs) from WT and Fyn^−/−^ mice

mESCs were obtained from the inner cell mass of blastocysts isolated from 3.5 days post coitum mice^47^. Briefly, a feeder layer of mouse embryonic fibroblasts (MEFs, 1×10^6^) was plated in tissue culture dishes two days before blastocyst isolation and treated with mitomycin C (10 µg/mL) for 2 h. Isolated blastocysts were washed with KnockOut^TM^ Dulbecco’s Modified Eagle Medium (DMEM), transferred aseptically to the feeder layer, and cultured with KnockOut^TM^ DMEM supplemented with 20 % knockout serum replacement and 1000 U/mL Leukemia Inducing Factor (LIF). After 48 h the medium was replaced, and the blastocysts were left to hatch for 7-10 days. Cells released from hatched blastocysts were transferred to a new feeder layer. Colonies generated from the previous step were transferred onto a new feeder layer. Pluripotency was confirmed through alkaline phosphatase (AP) staining and immunofluorescence of Sox2 and KLF4 (Supplementary Fig. 7A-D).

### Alkaline phosphatase (AP) staining

To perform AP staining, mESCs were fixed with 4 % paraformaldehyde in phosphate buffered saline (PBS) for 2 min at room temperature. AP staining was performed using an Alkaline Phosphatase Detection Kit following the manufacturer’s protocol (MilliporeSigma).

### mESCs culture

The mESCs E14Tg2a line was maintained in DMEM supplemented with 15 % heat-inactivated fetal bovine serum (FBS), 0.055 mM β-mercaptoethanol, 2 mM L-glutamine, 0.1 mM minimum essential medium non-essential amino acids, and 1,000 U/mL LIF.

### RNA interference of Fyn

To generate Fyn knockdown cells, pLKO.1-sh-Fyn or pLKO.1-mock lentivirus plasmids were co-transfected with packaging vectors psPAX2 and pMD2.0G into HEK 293T cells using the iMFectin transfection reagent (GenDepot). The mouse lentiviral sh RNA vector sequence for Fyn was designed and confirmed in the University of Minnesota Genomic Center. Viral particles containing 4 μg/mL polybrene were transduced into E14TG2a cells at 24 h and 48 h time points. After transduction, cells were selected in the presence of 1 μg/mL puromycin for 7-10 days.

### Telomeric repeat amplification protocol (TRAP) assay

TRAP assay was used to determine telomerase activity according to the manufacturer’s recommendation (TRAPEZE® RT Telomerase Detection Kit S7710, MilliporeSigma). Briefly, cell lysates were resuspended into CHAPS lysis buffer at a concentration of 500 ng/µL. Relative telomerase activity was determined using a TSR8 standard curve following the protocol described by the manufacturer.

### Quantitative Fluorescent In Situ Hybridization (Q-FISH) and flow cytometry-FISH (Flow-FISH) assays

Q-FISH of metaphase chromosomes, interphase nuclei, and mouse tissues, was performed as previously described^48^. Briefly, cells were treated with 1 µg/mL colcemid (Cayman Chemicals) for 1 hour, hypotonically swelled with 75 mM KCl solution, and fixed with Carnoy’s solution. Embedded tissue samples were cut in 5 µM sections and deparaffinized using standard protocols. Telomere and centromere Q-FISH staining was performed using a TelC-Alexa-488, or TelG-Cy3 PNA telomere probe together with a CENPB-Cy3 centromere probe (PNA Bio Inc). Slides were counterstained with DAPI (Electron Microscopy Sci.) and visualized using a Nikon or a Zeiss LSM 900 confocal microscope. Telomere length was measured using an open-source software (Telometer; http://demarzolab.pathology.jhmi.edu/telometer/), as previously described^49^.

Flow-FISH analysis was performed as previously described^50^ with slight modifications. Briefly, 0.5 x 10^6^ cells were washed with PBS containing 0.1 % BSA and resuspended in 300 µL of hybridization buffer (70 % formamide, 20 mM Tris-HCl, pH 7.0, 1 % blocking reagent, and 1 µg/mL PNA probe) and incubated for 2 h at RT. Cells were washed twice with fixation buffer (60 % formamide, 10 mM Tris-HCl, pH 7.0, 0.1 % blocking reagent, 0.1 % Tween 20) at 40 °C, resuspended in PBS containing 0.1 % BSA and 0.05 µg/mL propidium iodide (PI) and analyzed using a BD-FACS (BD Aria III). TelC-Alexa-488 PNA probe and PI nuclear staining was used in all flow-FISH analysis. CRISPR-Cas9 transfected cells were stained with the TelG-Cy3 PNA probe and counterstained with DAPI.

### TeloSizer® assay

Telomere length was evaluated using the TeloSizer® assay (Genomic Vision, Bagneux, France) based on molecular combing technology. This assay measures physical telomere lengths in kilobase (kb), but also differentiates TTS (true telomere sequence) and ITS (interstitial telomere sequence).

Briefly, cells were harvested and embedded into agarose plugs using the Genomic Vision FiberPrep® kit (Genomic Vision, Bagneux, France). DNA extraction and combing were performed according to the EasyComb procedure (Genomic Vision, Bagneux, France). Coverslips were hybridized with a telomere-specific probe G-rich PNA probe Cy3-(TTAGGG)3 (Panagene, Daejeon, South Korea) followed by counterstaining with YOYO™-1 Iodide (Thermo Scientific, Bordeaux, France) according to the TeloSizer® assay procedure. Coverslips were scanned using the FiberVision® scanner. Telomere signals were automatically detected and measured using the FiberStudio® Easyscan software. TeloSizer® assay is based on molecular combing and has been shown to provide accurate telomere length measurements comparable to other telomere length analysis techniques^51^.

### Generation of DC-iPSCs from fibroblasts

DC fibroblasts were purchased from The Coriell Cell Repositories and maintained in MEM supplemented with 15 % FBS and 1 % non-essential amino acids. Induced pluripotent stem cells (iPSCs) were generated from DC fibroblasts by lentiviral transduction with polycistronic STEMCCA vector containing transfecting Yamanaka transcription factors Oct4, Klf4, Sox2, and c-Myc according to the manufacturer’s protocol (MilliporeSigma). Briefly, 1 x10^6^ DC fibroblasts were seeded onto gelatin-coated dishes and transduced with the STEMCCA lentivirus. Transduced cells were transferred to a plate containing 1 x 10^5^ feeder cells. iPSC colonies were picked 25–30 days post-transduction and maintained in feeder-free E8 media (StemCell Technologies) on a Matrigel (Corning). Pluripotency was confirmed by immunofluorescence and Western blotting using human iPSC specific markers (Supplementary Figs. 17A-B).

### q-PCR assay for telomere length measurement

Average telomere length was measured using a qPCR method previously described^52^. The single-copy genes *m36B4* and *h36B4,* and the multicopy gene *hAlu* were used as normalization references. A 20 μL PCR reaction contained 10 μL of 2X SYBR Green mix, 0.5 μL of each 10 μM forward and reverse primers (Supplementary Table 5), 4 μL DNase/RNase free water, and 5 μL genomic DNA at a concentration of 8 ng/μL. The qPCR was carried out in a Bio-Rad thermocycler (CFX96) with reaction conditions of 95 °C for 10 min followed by 40 cycles of denaturation at 95 °C for 15 s, 60 °C annealing for 30 s, 72 °C extension and data collection for 30 sec. The CFX manager software was used to generate standard curves. The average telomere length is expressed as the telomere to single copy gene (T/S) ratio.

### Protein extraction, Western blotting, and Immunohistochemistry (IHC)

Cells were lysed with NP-40 lysis buffer (50 mM Tris-HCl, pH 7.6, 150 mM NaCl, 5 mM EDTA, 1 % Nonidet P-40 with a protease inhibitor cocktail). Nuclear and Cytoplasmic fractions were obtained using the NE-PER cytoplasmic and nuclear protein extraction kit (ThermoFisher Scientific). A phosphatase inhibitor cocktail was used for phosphoprotein analysis, and N-ethylmaleimide (NEM) was used for SUMO-protein analysis. Proteins were resolved in SDS-PAGE and transferred to polyvinylidene difluoride (PVDF) membranes. The membranes were blocked with 5 % milk, and target proteins were detected with specific antibodies and visualized by chemiluminescence.

IHC was performed using a standard protocol. Paraffin-embedded tissues were subjected to deparaffination and antibody retrieval. Tissue sections were blocked with 10 % goat-serum followed by overnight incubation with a primary antibody at 4 °C. Slides were incubated with a secondary antibody and stained with 3,3′-Diaminobenzidine (DAB). For IF, a fluorochrome-conjugated secondary antibody was used. Images were quantified using ImageJ2 software^53^.

### Immunoprecipitation

A total amount of 1 mg of cell lysates in a volume of 500 µL were immunoprecipitated with 2 µg of target antibody and samples were rotated overnight at 4 °C. 40 µL of protein A/G Sepharose beads (GenDepot) were added to each sample and rotated 1 h at 4 °C. Beads were washed 3 times with lysis buffer, and the supernatant fraction was analyzed by Western blotting.

### Chromatin immunoprecipitation assay

Stable sh-mock and sh-Fyn E14 cells were seeded in 10 cm^2^ plates. Chromatin immunoprecipitation (ChIP) was performed using the One-Day Chromatin Immunoprecipitation Kit (Magna ChIP G, MilliporeSigma) according to the manufacturer’s protocol. Chromatin samples were immunoprecipitated with anti-Menin antibody overnight at 4 LJ. The DNA fractions for TERT promoter was analyzed by qPCR.

### Protein expression, purification, and in vitro kinase assay

WT and mutant *MEN1* (Y603F) were cloned into a pGEX-5X1 vector upstream a C-terminal 6×His tag. Menin WT and mutant proteins were expressed in *E. coli* BL21(DE3) pLysS cells (Promega). Cells were grown in Lysogeny broth to an OD 600 of 0.5-0.7 and protein expression was induced with 0.5 mM IPTG for 20 h at 16 °C. Cells suspended in 200 mL were pelleted at 6000 × g and resuspended in 10 mL lysis buffer (50 mM HEPES pH 8.0, 500 mM NaCl, 20 mM imidazole, 10 % glycerol, 2 mM TCEP, 1 % Tween-20, and 1X protease inhibitor). Cell suspensions were incubated with lysozyme for 30 min and disrupted by sonication for 10 min. Cells were centrifuged for 30 min at 20000 × g, and lysates were equilibrated with Ni-NTA agarose beads (ThermoFisher Scientific) for 2 h at 4 °C. Samples were centrifuged at 120 × g for 3 min to remove the supernatant fraction. Beads were washed with 10 mL buffer 1 (50 mM HEPES pH 8.0, 500 mM NaCl, 30 mM imidazole, 10 % glycerol, 0.5 % Tween-20, and 2 mM DTT), and 2X with 10 mL buffer 2 (50 mM HEPES pH 8.0, 500 mM NaCl, 40 mM imidazole, 10 % glycerol, 0.5 % Tween-20, and 2 mM DTT). Proteins were eluted with three washes of 0.1 mL elution buffer (50 mM HEPES pH 8.0, 150 mM NaCl, and 300 mM imidazole) and transferred to a kinase buffer (50 mM Tris pH 8.0, and 100 mM NaCl) using Zeba Spin Desalting columns (ThermoFisher Scientific). Proteins were separated by SDS-PAGE and stained with Coomassie blue. The protein concentrations were quantified using a Nanodrop (ThermoFisher Scientific).

For radioactive *in vitro* kinase assay, 100 ng of Fyn protein (MilliporeSigma, Cat # 14-441) was incubated with or without 1 µg of recombinant Menin protein, either commercially purchased (Novus Biologicals, Cat # H00004221-P01) or produced in the laboratory. WT or mutant Menin were incubated for 30 min in a 30 °C water bath. Proteins were incubated with 10 μCi [^γ32^P] ATP in kinase buffer (20 mM HEPES (pH 7.4), 1 mM dithiothreitol, 10 mM MgCl_2_, and 10 mM MnCl_2_). The incorporated radioactivity was measured by autoradiography.

For the non-radioactive kinase assay, 100 ng of commercially purchased Fyn protein (MilliporeSigma) was incubated with or without in house synthesized 1 µg Menin recombinant protein (WT or mutant Menin) for 30 min in a 30 °C water bath. Proteins were incubated in the presence of 200 µM ATP in kinase buffer (20 mM HEPES (pH 7.4), 1 mM dithiothreitol, 10 mM MgCl_2_, and 10 mM MnCl_2_). Immunoblots measured the incorporated ATP phosphorylation to Menin using a pY603-Menin antibody. Lambda protein phosphatase was used to validate the specificity of the pY603-Menin antibody. To stop the Fyn kinase activity, the *in vitro* kinase assay reaction mix was incubated for 30 min in a 30 °C water bath with 400 U of Lambda protein phosphatase in the presence of 10 mM MnCl_2_ in 1x NEBuffer for Protein MetalloPhosphatases. To counteract the effect of Lambda protein phosphatase, we added a 1X phosphatase inhibitor cocktail to the reaction mix to recover the phospho signal. The *in vitro* kinase assay reaction mixtures were separated into 8 % polyacrylamide gels and blotted against a pY603-Menin. Membranes were stripped and reblotted with Menin and Fyn antibodies.

For *in vivo* kinase assay, Xpress-Menin and HA-Fyn were over-expressed in 293T cells. Xpress-Menin was immunoprecipitated with Xpress antibodies and the Lambda phosphatase assay was performed with or without 1× phosphatase inhibitors as previously described. Reaction mixtures were separated into 8 % PAGE and blotted with pY603-Menin, Menin or Fyn antibodies. *In vivo* kinase assays were also performed in endogenously expressed Menin protein. Briefly, endogenous Menin was immunoprecipitated with an anti-Menin antibody (Invitrogen) and Lambda phosphatase assay was performed as previously described. Reaction mixtures were separated into 8 % polyacrylamide gels and blotted against a pY603-Menin or Menin antibody.

### Site-directed mutagenesis

Site-directed mutagenesis was performed using QuikChange II Site-Directed Mutagenesis Kit (Agilent) according to the manufacturer’s protocol. Primers for mutagenesis were designed using online tools provided by Agilent. WT MEN1 plasmid was used as a template. The mutations were confirmed by sequencing.

### Identification of phosphorylation sites on Fyn-phosphorylated Menin

*An in vitro* kinase assay was performed as discussed above, replacing [^γ32^P]ATP with unlabelled 200 µM ATP. The ABSciex TripleTOF 5600 system (SCIEX) coupled with Eksigent 1D+ Nano LC system (SCIEX) was used to identify Menin’s phosphorylated sites. Nano LC of enzymatic peptides was performed with the Eksigent 1D+ Nano LC equipped with the cHiPLC nanoflex system (SCIEX). Peptides were loaded onto the column and then eluted with a linear gradient of 5 – 40 % binary solvent B1 for 30 min at a flow rate of 0.3 mL/min. Mass spectrometry analysis of peptides was performed using the ABSciex TripleTOF 5600 system. Analysis was performed using the NanoSpray III source (SCIEX). The mass spectrometry was calibrated by the acquisition of [Glu1] fibrinopeptide (25 pmol/mL). The raw data were processed and searched with ProteinPilot^TM^ software (version 4.0) using the Paragon algorithm. Proteins were identified by searching the UniProtKB human database and filtered at an R95 % confidence cut-off. Peptides for phosphorylated Men1 were identified at a 1 % false discovery rate.

### Small ubiquitin-like modifier (SUMOylation) assay

We performed an *in vitro* SUMOylation assay with endogenously-expressed or overexpressed proteins. Briefly, HEK 293T cells were transfected with pcDNA3.1-Xpress-*MEN1* (WT or Y603F, Y603D, K493R, K609R mutants) and pcDNA3-HA-*SUMO-1* using iMFectin. 24 h after transfection, cells were lysed in NP-40 buffer. SUMOylated-Menin proteins were immunoprecipitated with Dynabeads™ Protein A beads (Invitrogen) coupled to Xpress tags or Protein A/G-Agarose (GenDEPOT) coupled to a Menin antibody and visualized with HA or SUMO1 antibodies. In other experiments we also pulldown Dynabeads™ Protein A beads (Invitrogen) coupled to HA or Menin antibody and visualized Menin-SUMOylation by blotting against Xpress and HA respectively. We analyzed SUMOylation of Menin by using the EpiQuik^TM^ In Vivo Universal Protein Sumoylation Assay Kit (EPIGENTEK). Briefly, nuclear lysates from mESCs were prepared using the NE-PER cytoplasmic and nuclear protein extraction kit (ThermoFisher Scientific). SUMOylated proteins were pulldown using Dynabeads™ Protein A beads and 10 µg of pulldown protein was used for the assay. SUMOylation intensity was calculated against a standard curve prepared from recombinant SUMO protein according to the manufacturer’s protocol.

### RNA-immunoprecipitation (RNA-IP) and RNP-immunoprecipitation (RNP-IP)

RNA-IP was performed according to Tang *et al*., 2018^54^. Briefly, cDNA was used as a template to amplify hTR or mTR. The 5′ primers contained the T7 promoter sequence 5′-CCAAGCTTCTAATACGACTCACTATAGGGAGA-3′. PCR-amplified DNA was used as a template to transcribe biotinylated RNA using T7 RNA polymerase and biotinylated-UTP. One µg of purified biotinylated transcripts was incubated with 200 μg of nuclear extracts for 60 min at 4 °C. Complexes were isolated with streptavidin-conjugated agarose beads (MilliporeSigma), and the pull-down material was analyzed by Western blotting.

For RNP IP assays, cells were exposed to UVC (400 mJ/cm^2^), and lysates were prepared in IP buffer (10 mM Hepes, pH 7.4, 50 mM β-glycerophosphate, 1 % Triton X-100, 10 % glycerol, 2 mM EDTA, 2 mM EGTA, 10 mM NaF, 1 mM DTT, protease inhibitor cocktail, NEM, RNase inhibitor) and immunoprecipitated with anti-Menin or anti-Xpress antibodies overnight. The complexes were washed twice with stringent buffer (100 mM Tris-HCl, pH 7.4, 500 mM LiCl, 0.1 % Triton X-100, and 1 mM DTT, protease inhibitor cocktail, N-ethylmaleimide, and RNase inhibitor) and twice with IP buffer. The RNA in RNP IP was assessed by qRT-PCR analysis and normalized with the copy number of the *U6* gene^54^.

### Mass spectrometry analysis of RNA-IP

A proteome-based approach was used for determining the protein targets associated with TR after transfection with *SUMO1* and *MEN1* Y603F or Y603D. First, HEK 293T cells were transfected with *MEN1* mutants (Y603F or Y603D) and HA-tagged *SUMO-1* for 24 h. Nuclear lysates from the transfected cells were subjected to RNA-IP with TR. Differential protein expression across the samples was screened by conducting MS/MSALL with SWATHTM acquisition. The raw data were collected through Information Dependent Acquisition (IDA) and SWATHTM acquisition using ABSciex TripleTOF^TM^ 5600 with Eksigent 1D+ nano LC (nanoLC-MS/MS instrument). Protein identification, MS peak extraction, and statistical analysis were performed with ProeinPilot^TM^ (version 4.5), PeakView^TM^ (version 2.2) and MarkerView^TM^ (version 1.2), respectively. Bioinformatics analyses were conducted with the BioVenn program and displayed in a Venn-diagram of differential expressed proteins^55^. Protein classification was performed using the String software (https://string-db.org) and Graphpad prism 8.0 was used to make the pie diagram.

### Cycloheximide Chase Assay

Cycloheximide (CHX) chase assay was performed in transfected HEK 293T cells using treatment of 50 µg/mL CHX (Cayman Chemical) at different time points. Briefly, 5×10^5^ cells were plated in a 6 well plate and transfected with Y603F*-MEN1* and Y603D-*MEN1* with or without *SUMO1.* Transfected media was replaced by fresh media containing 50 µg/mL CHX and treated for 0, 4 and 8 h. Lysate were prepared using M-PER^TM^ mammalian protein isolation buffer (ThermoScientific).

### Generation of Fyn knockout DC-iPS cells by CRISPR-Cas9

CRISPR-Cas9-mediated genome editing was performed transfecting DC fibroblasts with Fyn and control Double Nickase human Plasmids (Santa Cruz Biotechnology, Supplementary Table 6). according to the manufacturer’s protocol. Immunoblots confirmed gene knockdown efficiency.

### pY603-Menin antibody validation

The specificity of custom purified pTyr603-menin (pY603-menin) rabbit polyclonal antibody (Abclonal) was confirmed using a Lambda phosphatase assay (New England BioLabs).

### Immunofluorescence-FISH (IF-FISH)

Unsynchronized cells were fixed on culture slides with 4 % paraformaldehyde at room temperature for 10 min, followed by permeabilization with 0.05 % Triton-X100 for 10 min. Slides were blocked in 5 % serum for 1 h, incubated overnight with a primary antibody, and incubated with a fluorochrome-tagged secondary antibody for 1 h. The same slides were subjected to FISH assay using telomeric PNA probes as described above.

FISH of TR was performed using the Single-Molecule Inexpensive FISH (smiFISH) method with slight modifications^56^. Briefly, FLAP oligos were annealed to 15 human telomeric RNA (hTR) probes (Supplementary Table 7) using a PCR machine with a denaturation step at 85 °C for 3 mins, and annealing at 65 °C for 3 mins and 25 °C for 5 mins. The probes used in this experiment recognize human TR RNA as well as mouse TR RNA due to high sequence similarity. About, 1× 10^3^ cells were grown onto 0.5 % gelatin-coated 4/8 well glass chambered slides (MilliporeSigma). Cells were fixed with 4 % paraformaldehyde for 20 min, rinsed twice with PBS, and permeabilized with 0.5 % Triton-X-100 in PBS for 5 min. After permeabilization, cells were washed twice with PBS followed by one wash with 1x SSC 15 % formamide buffer, and incubated with the same buffer for an additional 15 min. Cells were then covered with smiFISH hybridization buffer (1x SSC, 15 % formamide, 4.5 mg/mL BSA, 10.6 % dextran sulfate, 2 mM VRC, 0.425 mg/mL tRNA, 0.08 µM FLAP-annealed probe set) and incubated at 37 °C overnight in a humidified chamber protected from light.

Slides were washed twice with 1X SSC, 15 % formamide at 37 °C for 30 mins, followed by rinsing twice in PBS. The slides were then blocked with IF antibody buffer (0.1 % Triton-X-100 and 2 % BSA in PBS) for 2 h and incubated with the primary antibody dissolved in the same buffer at 4 °C overnight. Slides were washed three times with 0.1 % Triton-X-100 in PBS and twice with the IF antibody buffer. Slides were incubated with Alexa fluor secondary antibodies resuspended in the IF antibody buffer for 1 h at room temperature. Slides were washed twice with IF antibody buffer and three times with 0.1 % Triton-X-100 in PBS and covered with DAPI with antifade.

For IF double FISH, TR FISH was performed followed by telomere FISH using a PNA probe as we previously described. IF was performed on the same slides.

### Statistical analysis

Data were analyzed either by student’s t-test or by one-way analysis of variance (ANOVA) followed by Tukey multiple comparisons test using the GraphPad Prism software (Ver 9.0). The frequency of distribution and detailed statistical analysis was performed using SPSS software (Ver 29.0). PCR data were obtained in at least three replicates. Western blot and ELISA quantitation was performed with at least three independent experiments. Immunofluorescent and FISH data were analyzed in at least 10 micrographs, unless it is indicated otherwise. Each experiment was performed at least 3 times unless otherwise indicated. A probability of *p* < 0.05 was considered statistically significant. ^∗^*p* ≤ 0.05, ^∗∗^*p* ≤ 0.01, and ^∗∗∗^*p* ≤ 0.001

## Acknowledgements

The authors thank Genomic Vision’s ‘TeloSizer® Triathlon Program’ (Genomic Vision, Bagneux, France) and their help for Molecular Combing Assay. The authors are grateful to Tara Adams and Teri Bishop Johnson for their help to handle animal colonies, and Josh Monte and Todd Schuster for their help in Flow-FISH analysis. NIH grant R01-GM-143428 supported R.C-G. and W.C. An NIH/NCI grant R21-CA259630 and the National Scleroderma Foundation supported R.C-G. This work was supported by The Hormel Foundation.

