## Supplemental file for "Fyn-mediated phosphorylation of Menin disrupts telomere maintenance in stem cells"

### Supplementary Figure Legends

**Supplementary Fig. 1. Fyn inhibition slows telomere erosion in colon tissue of mice**

Related to Fig. 1. (A), Representative images of Q-FISH analysis using telomere and centromere probes on colon tissue of 43-week-old wild type and $Fyn^{-/-}$ female mice, showing that Fyn knockout mice have significantly higher telomere fluorescence (green dots). The scale bar is 20 µm. A pseudo color grey was used replacing blue for nucleus.

(B, C) Frequency of distribution and detailed statistics of mean telomere fluorescence quantified from the above figure using Telometer in colon tissue from WT and $Fyn^{-/-}$ mice (n=1947 and 1805 reads, respectively). Q1, quartile 1; Q2, Quartile 2; Q3, quartile 3.

(D-F) Differences in mean telomere fluorescence (D), mean centromere fluorescence (E), and the ratio of telomere/centromere fluorescence (F) in WT and $Fyn^{-/-}$ mice. Centromere FISH signals were used as a staining control. Data are presented as fluorescent means ± SD from a two-tailed t-test (*** = $p < 0.001$, **** = $p < 0.0001$). A probability of $p < 0.05$ was considered statistically significant.

**Supplementary Fig. 2. Fyn inhibition slows telomere erosion in skin tissue of mice**

Related to Fig. 1. (A), Representative images of Q-FISH analysis using telomere and centromere probes on skin tissue of 43-week-old wild type and $Fyn^{-/-}$ female mice, showing that Fyn knockout mice have significantly higher telomere fluorescence (green dots). The scale bar is 20 µm. A pseudo color grey was used replacing blue for nucleus.

(B, C) Frequency of distribution and detailed statistics of mean telomere fluorescence quantified from the above figure using Telometer in colon tissue from WT and $Fyn^{-/-}$ mice (n=1947 and 1805 reads, respectively). Q1, quartile 1; Q2, Quartile 2; Q3, quartile 3.

(D-F) Differences in mean telomere fluorescence (D), mean centromere fluorescence (E), and the ratio of telomere/centromere fluorescence (F) in WT and $Fyn^{-/-}$ mice. Centromere FISH signals were used as a staining control. Data are presented as fluorescent means ± SD from a two-tailed t-test (*** = $p < 0.001$, **** = $p < 0.0001$). A probability of $p < 0.05$ was considered statistically significant.

**Supplementary Fig. 3. Fyn deletion or inhibition slows telomere erosion in stem cells from mice**

Related to Fig. 1. (A, B) Representative images of morphology (A) and AP staining (B) of mESCs isolated from WT and $Fyn^{-/-}$ mice. The scale bars are 50 and 20 μm, respectively.

Related to Fig. 1, (C, D) IF analysis of stem cell markers KLF4 and SOX2 in mESCs isolated from WT and $Fyn^{-/-}$ mice. The scale bar is 10 μm.

Related to Fig. 1. Representative images of interphase nuclei Q-FISH analysis on wild type vs $Fyn^{-/-}$ mESCs at passage P12 (E, top) and sh-Mock vs sh-Fyn E14 cells at passage 18 (E, bottom) The scale bar is 5 μm. Quantification of fluorescent signal using the Telometer program in WT vs $Fyn^{-/-}$ mESCs (F) and in sh-Mock vs sh-Fyn E14 cells (G), showing higher telomere fluorescence intensity in Fyn deficient conditions. n = 20 interphase nuclei.

Data are presented as means ± SD from a two-tailed t-test (** = $p < 0.01$, *** = $p < 0.001$). A probability of $p < 0.05$ was considered statistically significant.

**Supplementary Fig. 4. Fyn deletion or inhibition slows telomere erosion in stem cells**

(A, B, C, D) Telomere erosion is stabilized in $Fyn^{-/-}$ vs WT mESCs (A, B) and in sh-Fyn vs sh-Mock-transfected cells (C, D) as determined by telomere measurement by Flow-FISH analysis (A, C) and T/S ratio calculated by quantitative PCR (qRT-PCR), respectively (B, D). T/S ratios were calculated relative to the single-copy gene 36B4. Data are presented as means ± SD from a two-tailed t-test (* = $p < 0.05$, ** = $p < 0.01$).

Q-FISH analysis of sh-Mock, sh-Fyn and sh-Fyn/Fyn over-expression in E14 cells, showing that overexpression of the *Fyn* gene in sh-Fyn cells promotes telomere erosion (E). Western blotting analysis confirmed Fyn knockdown (F) and over-expression status (G). β-actin and tubulin act as loading controls. Molecular weight markers are shown at the right. Mean telomere fluorescence was calculated using Telometer from panel E (H).

Data are presented as means ± SD from one-way ANOVA followed by Tukey's multiple comparison test (**** =$p < 0.0001$).

A probability of $p < 0.05$ was considered statistically significant.

**Supplementary Fig. 5. Fyn deletion improves telomere length in mice**

(A) Representative image of genotyping of wild type $Fyn^{+/+}$, heterozygous $^{-/+}$, and $Fyn^{-/-}$ mice.

(B) IHC analysis of Fyn in colon or skin tissue of female and male wild type and $Fyn^{-/-}$ mice. The scale bar is 20 µm.

(C-F) Quantitation of IHC images using imageJ shows reduced expression of Fyn in colon (C) and skin tissue (D) of female $Fyn^{-/-}$ mice or colon (E) and skin tissue (F) of male $Fyn^{-/-}$ mice. Data are presented as means ± SD. An unpaired student's t-test was used for statistical analysis. A probability of p < 0.05 was considered statistically significant (**** = $p < 0.0001$).

**Supplementary Fig. 6. Fyn inhibition slows telomere erosion in 97 weeks mouse tissues**

(A, B) Representative images of female and male colon (A) and skin tissues (B) from 97-week-old WT and $Fyn^{-/-}$ mice used in Q-FISH analysis. Scale bar is 20 µm. The mean Telomere fluorescence indexes shown in a graph plot at the right of the figures were quantitated using the Telometer program (n = 100 nuclei). Arrows indicate telomeres. Data are presented as mean ± SD from a two-tailed t-test (**** = $p < 0.0001$). A probability of $p < 0.05$ was considered statistically significant.

**Supplementary Fig. 7. Fyn does not inhibit telomerase activity *in vitro.***

Western blotting of Fyn and TERT in (A) WT and $Fyn^{-/-}$ mESCs and sh-Mock and sh-Fyn cells (B). β-actin served as a loading control. Densitometry analysis by ImageJ shows no significant difference in TERT expression. Molecular weight markers are shown at the right.

Telomerase enzyme activity (TRAP assay fold change) was not significantly different between WT and $Fyn^{-/-}$ mESCs (C), between sh-Mock and sh-Fyn E14 cells (D), in colon of 43-weeks-old WT vs $Fyn^{-/-}$ mice (E), or sh-Mock and sh-Fyn 293T cells at passage 10 (F).

Data are presented as means ± SD from a two-tailed t-test. A probability of $p < 0.05$ was considered statistically significant. ns = not significant.

**Supplementary Fig. 8. Fyn interacts with Menin**

Immunoprecipitation using an anti-Fyn antibody followed by Western blotting using proteins that influence the telomere pathway showing only Menin interacting with Fyn (scattered box). Arrows indicate the respective protein band sizes.

**Supplementary Fig. 9. Fyn phosphorylates Menin at tyrosine 603**

Related to Fig. 3, (A, B, C) Tandem mass spectrometry spectra of the Menin site phosphorylated by Fyn. The precursor mass (m/z) of 611.7975 was matched to the triply charged peptide, DpYTLSFLKR, indicating that Y603 is phosphorylated. The b2 ion at 359.06 and b3 ion at 460.11 indicate that Y603 is phosphorylated.

**Supplementary Fig. 10. Menin does not bind to TERT promoter or with TERT in mESCs**

(A) Menin occupancy at the TERT promoter in sh-Mock and sh-Fyn E14 cells after chromatin immunoprecipitation (ChIP) of Menin with an anti-Menin antibody and amplification of TERT by qPCR. Data are presented as the fold enrichment mean ± SD of the TERT promoter amplification. An unpaired student's t-test was used for statistical analysis. ns = not significant.

(B) Immunoprecipitation of TERT using an anti-Menin antibody in Sh-Mock and Sh-Fyn E14 cells followed by Western blotting using an anti-Menin antibody shows Menin and TERT are not binding partners.

**Supplementary Fig. 11. Menin is not SUMO-modified by SUMO2/3/4. Fyn-deficient cells have decreased Menin-SUMO1 modification**

(A) Prediction results of Menin SUMOylation sites from the GPS-SUMO open-source program.

(B) Immunoprecipitation of endogenous Menin and immunoblot against SUMO2/3/4 in sh-Mock and sh-Fyn transfected E14 cells, showing the lack of interaction between Menin and SUMO2/3/4.

(C) Menin SUMO1 modification is decreased in Fyn-deficient cells as shown by ELISA in nuclear lysates of WT and $Fyn^{-/-}$ mESCs or sh-Mock and sh-Fyn E14 cells. Data are presented as SUMOylation intensity means ± SD from a two-tailed t-test (*** = $p < 0.001$). A probability of $p$

< 0.05 was considered statistically significant.

**Supplementary Fig. 12. Menin Phosphorylation and SUMOylation promote TR binding through improved protein stability**

Related to Fig. 6, (A) Phosphorylation and SUMOylation at the C-terminal site of Menin contribute for Menin-TR binding. RNA pulldown assay using *in vitro* transcribed TR and overexpressed mutant (Y603F, Y603D, K609R) Menin with SUMO1 indicates that 50 times lesser protein of Y603D Menin with SUMO1 interacts with TR very strongly compared to Y603F and K609R Menin with SUMO1.

(B, C) Cycloheximide chase assay showing significantly higher protein degradation in Y603F-Menin with or without SUMO1 compared to Y603D-Menin with or without SUMO1. Y603F and Y603D transfected HEK 293T cells co-transfected with SUMO1 were treated with 50μg/mL cycloheximide for indicated time points. Xpress-tagged Menin (B) and Menin (C) protein expression were analyzed at indicated time points. Differential exposure times show very strong expression of only Xpress-Menin in Y603D and Y603D+SUMO1 transfected cell in 1 min exposure. Y603F and Y603F+SUMO1 were visible after 10 mins. Blotting with anti-Menin antibody shows Menin band at short exposure. Menin was analyzed with an Xpress antibody that binds the C-terminal Xpress Tag of the protein or a Menin antibody that binds the central region of Menin.

**Supplementary Fig. 13. SUMO1-modified Menin prevents TR's association with Dyskerin**

Related to Fig. 7. Representative IF-FISH images show Coilin and TR binding in Y603F-MEN1+SUMO1, K609R-MEN1+SUMO1, and Y603D-MEN1+SUMO1 transfected 293T cells.

The scale bar is 5 μm. Arrows indicate co-localization. A pseudo color yellow was used replacing magenta for nucleus.

**Supplementary Fig. 14. Generation of DC-iPSC from DC- fibroblasts**

(A) Immunofluorescence analysis of SSEA-4 and TRA-1-60 protein expression, showing positive staining for transduction pluripotency markers in DC iPSCs. The scale bar is 100 μm.

(B) Western blotting analysis of pluripotency markers Nanog and KLF4 showing expression in DC iPSCs, but not in DC fibroblasts. Molecular weight markers are shown at the right.

**Table S1.** Candidate Fyn substrate's list for probable telomere length regulator

| Transcription factor | Activator/Repressor | Reference |
|---|---|---|
| Stat1 | Unknown | |
| Stat3 | Activator | Ref. 19 |
| Stat5 | Activator | |
| c-Myc | Activator | |
| KLF2 | Repressor | |
| KLF7 | Unknown | |
| KLF15 | Unknown | |
| Menin | Repressor | Ref. 19 |
| E2F1 | Repressor | |
| SP3 | Repressor | |
| WT1 | Repressor | |
| MAD1 | Repressor | |

**Table S2.** Differential TERC-bound proteins detected in Mass-spectrometry analysis using RNA-IP products of Y603F-*MEN1+SUMO-1* and Y603D-*MEN1+SUMO-1* transfected 293T cells.

**Y603F-Menin+SUMO-1 -TERC binding proteins:**

| Accession # | Name |
|---|---|
| sp\|Q00839\|HNRPU_HUMAN | Heterogeneous nuclear ribonucleoprotein U |
| sp\|P22626\|ROA2_HUMAN | Heterogeneous nuclear ribonucleoproteins A2/B1 |
| sp\|Q9NR30\|DDX21_HUMAN | Nucleolar RNA helicase 2 |
| sp\|O43390\|HNRPR_HUMAN | Heterogeneous nuclear ribonucleoprotein R |
| sp\|P16402\|H13_HUMAN | Histone H1.3 |
| sp\|Q99729-3\|ROAA_HUMAN | Isoform 3 of Heterogeneous nuclear ribonucleoprotein A/B |
| sp\|P09874\|PARP1_HUMAN | Poly [ADP-ribose] polymerase 1 |
| sp\|P07910-2\|HNRPC_HUMAN | Isoform C1 of Heterogeneous nuclear ribonucleoproteins C1/C2 |
| sp\|P52272\|HNRPM_HUMAN | Heterogeneous nuclear ribonucleoprotein M |
| sp\|Q92841\|DDX17_HUMAN | Probable ATP-dependent RNA helicase DDX17 |
| sp\|O94776\|MTA2_HUMAN | Metastasis-associated protein MTA2 |
| sp\|Q9Y3I0\|RTCB_HUMAN | tRNA-splicing ligase RtcB homolog |
| sp\|P98179\|RBM3_HUMAN | RNA-binding protein 3 |
| sp\|Q0P5I0\|MEN1_BOVIN | Menin |
| sp\|P29692\|EF1D_HUMAN | Elongation factor 1-delta |
| sp\|Q8N684\|CPSF7_HUMAN | Cleavage and polyadenylation specificity factor subunit 7 |
| sp\|P12956\|XRCC6_HUMAN | X-ray repair cross-complementing protein 6 |
| sp\|Q6ZRS2\|SRCAP_HUMAN | Helicase SRCAP |
| sp\|P55265\|DSRAD_HUMAN | Double-stranded RNA-specific adenosine deaminase |
| sp\|P26358\|DNMT1_HUMAN | DNA (cytosine-5)-methyltransferase 1 |

| | |
|---|---|
| sp\|Q13263\|TIF1B_HUMAN | Transcription intermediary factor 1-beta |
| sp\|Q969G3\|SMCE1_HUMAN | SWI/SNF-related matrix-associated actin-dependent regulator of chromatin subfamily E member 1 |
| sp\|Q1KMD3\|HNRL2_HUMAN | Heterogeneous nuclear ribonucleoprotein U-like protein 2 |
| sp\|Q92922\|SMRC1_HUMAN | SWI/SNF complex subunit SMARCC1 |
| sp\|O00571\|DDX3X_HUMAN | ATP-dependent RNA helicase DDX3X |
| sp\|Q9Y3C1\|NOP16_HUMAN | Nucleolar protein 16 |
| sp\|P24534\|EF1B_HUMAN | Elongation factor 1-beta |
| sp\|Q9Y4C8\|RBM19_HUMAN | Probable RNA-binding protein 19 |
| sp\|P07814\|SYEP_HUMAN | Bifunctional glutamate/proline--tRNA ligase |
| sp\|O60506\|HNRPQ_HUMAN | Heterogeneous nuclear ribonucleoprotein Q |
| sp\|Q8TAQ2\|SMRC2_HUMAN | SWI/SNF complex subunit SMARCC2 |
| sp\|Q9BXP5\|SRRT_HUMAN | Serrate RNA effector molecule homolog |
| sp\|Q5SRE5\|NU188_HUMAN | Nucleoporin NUP188 homolog |
| sp\|Q9H6R4\|NOL6_HUMAN | Nucleolar protein 6 |
| sp\|Q9NY12\|GAR1_HUMAN | H/ACA ribonucleoprotein complex subunit 1 |
| sp\|Q9Y2L1\|RRP44_HUMAN | Exosome complex exonuclease RRP44 |
| sp\|Q9P2I0\|CPSF2_HUMAN | Cleavage and polyadenylation specificity factor subunit 2 |
| sp\|Q9Y2X3\|NOP58_HUMAN | Nucleolar protein 58 |
| sp\|Q14980\|NUMA1_HUMAN | Nuclear mitotic apparatus protein 1 |
| sp\|Q14690\|RRP5_HUMAN | Protein RRP5 homolog |
| sp\|Q96I24\|FUBP3_HUMAN | Far upstream element-binding protein 3 |
| sp\|Q9UMS4\|PRP19_HUMAN | Pre-mRNA-processing factor 19 |
| sp\|P31942\|HNRH3_HUMAN | Heterogeneous nuclear ribonucleoprotein H3 |
| sp\|O95983\|MBD3_HUMAN | Methyl-CpG-binding domain protein 3 |
| sp\|Q9H5H4\|ZN768_HUMAN | Zinc finger protein 768 |
| sp\|Q86YP4\|P66A_HUMAN | Transcriptional repressor p66-alpha |
| sp\|Q13243\|SRSF5_HUMAN | Serine/arginine-rich splicing factor 5 |
| sp\|Q12906\|ILF3_HUMAN | Interleukin enhancer-binding factor 3 |
| sp\|O60832\|DKC1_HUMAN | H/ACA ribonucleoprotein complex subunit 4 |
| **Y603D+ SUMO-1-TERC binding proteins:** | |
| Accession # | Name |
| sp\|Q00839\|HNRPU_HUMAN | Heterogeneous nuclear ribonucleoprotein U |
| sp\|O43390\|HNRPR_HUMAN | Heterogeneous nuclear ribonucleoprotein R |
| sp\|Q99729-3\|ROAA_HUMAN | Isoform 3 of Heterogeneous nuclear ribonucleoprotein A/B |
| sp\|P09874\|PARP1_HUMAN | Poly [ADP-ribose] polymerase 1 |
| sp\|P52272\|HNRPM_HUMAN | Heterogeneous nuclear ribonucleoprotein M |
| sp\|Q9NR30\|DDX21_HUMAN | Nucleolar RNA helicase 2 |
| sp\|O60506\|HNRPQ_HUMAN | Heterogeneous nuclear ribonucleoprotein Q |
| sp\|Q9Y3I0\|RTCB_HUMAN | tRNA-splicing ligase RtcB homolog |
| sp\|P29692\|EF1D_HUMAN | Elongation factor 1-delta |
| sp\|Q14839\|CHD4_HUMAN | Chromodomain-helicase-DNA-binding protein 4 |
| sp\|Q99879\|H2B1M_HUMAN | Histone H2B type 1-M |

| | |
|---|---|
| sp|O00571|DDX3X_HUMAN | ATP-dependent RNA helicase DDX3X |
| sp|Q13263|TIF1B_HUMAN | Transcription intermediary factor 1-beta |
| sp|Q13435|SF3B2_HUMAN | Splicing factor 3B subunit 2 |
| sp|Q0P5I0|MEN1_BOVIN | Menin |
| sp|Q8N684|CPSF7_HUMAN | Cleavage and polyadenylation specificity factor subunit 7 |
| sp|Q14974|IMB1_HUMAN | Importin subunit beta-1 |
| sp|P98179|RBM3_HUMAN | RNA-binding protein 3 |
| sp|Q92841|DDX17_HUMAN | Probable ATP-dependent RNA helicase DDX17 |
| sp|O95983|MBD3_HUMAN | Methyl-CpG-binding domain protein 3 |
| sp|P07814|SYEP_HUMAN | Bifunctional glutamate/proline--tRNA ligase |
| sp|Q9BXP5|SRRT_HUMAN | Serrate RNA effector molecule homolog |
| sp|Q10570|CPSF1_HUMAN | Cleavage and polyadenylation specificity factor subunit 1 |
| sp|Q6ZRS2|SRCAP_HUMAN | Helicase SRCAP |
| sp|Q8TDD1|DDX54_HUMAN | ATP-dependent RNA helicase DDX54 |
| sp|P12956|XRCC6_HUMAN | X-ray repair cross-complementing protein 6 |

**Table S3. List of primary antibodies used for Western blotting (WB) and immunoprecipitation (IP)**

| Antigen | WB dilutions | IP dilutions | Antibody name | Catalog number | Company |
|---|---|---|---|---|---|
| Fyn[1] | 1:1000 | 1:100 | Fyn (15), Mouse | sc-434 | Santa Cruz |
| Fyn[2] | 1:1000 | —— | Fyn, Rabbit | 4023 | Cell Signaling |
| β-actin | 1:1000 | —— | β-Actin (C4), Mouse | sc-47778 | Santa Cruz |
| α-Tubulin | 1:1000 | —— | α-Tubulin (6A204), Mouse | sc-69969 | Santa Cruz |
| Lamin B1 | 1:1000 | —— | Lamin B1 (B-10), Mouse | sc-374015 | Santa Cruz |
| Menin[1] | 1:1000 | —— | Menin (B9), Mouse | sc-374371 | Santa Cruz |
| Menin[2] | 1:1000 | 1:100 | Menin, Rabbit | PA5-85330 | ThermoFisher |
| p-Menin | 1:2000 | —— | p(Y609)-Menin, Rabbit | Custom | Abclonal |
| Xpress | 1:5000 | 1:1000 | Anti-Xpress, Mouse | 46-0528 | Invitrogen |
| SUMO1 | 1:1000 | —— | SUMO-1 (D-11), Mouse | sc-5308 | Santa Cruz |
| SUMO-2/3/4 | 1:2000 | —— | SUMO-2/3/4 (C-3), Mouse | sc-393144 | Santa Cruz |
| HA | 1:1000 | —— | Anti-HA-HRP | 1201381900 | Roche |
| Nanong | 1:1000 | —— | Nanong (5A10) | Sc-134218 | Santa Cruz |
| KLF4 | 1:1000 | —— | KLF4, Rabbit | PA5-18058 | ThermoFisher |
| FLAG | 1:5000 | —— | M1 anti-FLAG® | F7425 | Millipore-Sigma |
| HA | —— | 1:100 | Anti-HA.11 | 901501 | BioLegend |

**Table S4. List of primary antibodies used for immunofluorescence (IF) and immunohistochemistry (IHC)**

| Antigen | IF/IHC dilutions | Antibody name | Catalog number | Company |
|---|---|---|---|---|
| SSEA-4 | 1:100 | Anti-HumanSSEA-4, MC-813-70, PE | 60062PE | STEMCELL Technologies |
| TRA-1-60 | 1:100 | Anti-Human TRA-1-60, TRA-1-60R, Alexa Fluor® 488 | 60064AD | STEMCELL Technologies |
| Menin | 1:200 | Menin (D45B1 XP®), Rabbit | 6891 | Cell Signaling |
| Fyn | 1:50 | Fyn (15), Mouse | sc-434 | Santa Cruz |
| Dyskerin | 1:100 | DKC1, Rabbit | PA5-28922 | ThermoFisher |
| Coilin | 1:800 | Coilin (D2L3J) XP®, Rabbit | 14168 | Cell Signaling |
| SOX2 | 1:100 | Sox-2 (E-4), Mouse | sc-365823 | Santa Cruz |
| KLF4 | 1:100 | KLF4, Rabbit | PA5-18058 | ThermoFisher |

**Table S5. Primer list**

| Primer Name | Primer Sequences (5'-3') |
|---|---|
| Telomere, forward | CGGTTTGTTTGGGTTTGGGTTTGGGTTTGGGTTT GGGTT |
| Telomere, reverse | GGCTTGCCTTACCCTTACCCTTACCCTTACCCTTA CCCT |
| m36B4, forward | ACTGGTCTAGGACCCGAGAAG |
| m36B4, reverse | TCAATGGTGCCTCTGGAGATT |
| h364, forward | CAGCAAGTGGGAAGGTGTAATCC |
| h364, reverse | CCCATTCTATCATCAACGGGTACAA |
| hAlu, forward | GACCATCCCGGCTAAAACG |
| hAlu, reverse | CGGGTTCACGCCATTCTC |
| Wild type-C57bl/6-genotyping, forward | AGGCCCAAGTTGACTATCCA |
| Fyn$^{-/-}$-C57bl/6-genotyping, forward | GGGAGGATTGGGAAGACAAT |
| Common, reverse | GGCAGCCTGTCTCAAAAGTC |
| MEN1-wt, forward | ACCGCTAGCATGGGGCTGAAGGCCGCCCAGAAG |
| MEN1-wt, reverse | ATTAAGCTTTTACTTATCGTCATCGTCGTACAGATC GAGGCCTTTGCGCTG |
| MEN1-Y603F, forward | TGAGGAAAGACAGAATGAAGTCACTAGGGGTGGAC |
| MEN1-Y603F, reverse | GTCCACCCCTAGTGACTTCATTCTGTCTTTCCTCA |
| MEN1-Y603D, forward | GAGGAAAGACAGAATGTCGTCACTAGGGGTGGACA |
| MEN1-Y603D, reverse | TGTCCACCCCTAGTGACGACATTCTGTCTTTCCTC |
| MEN1-K498R, forward | CTCCTCTGGCCTGGACTCCCGCCG |
| MEN1- K498R, reverse | CGGCGGGAGTCCAGGCCAGAGGAG |

| | |
|---|---|
| MEN1-K609R, forward | TGCGCTGCCGCCTGAGGAAAGACAGAATGTAGT |
| MEN1- K609R, reverse | ACTACATTCTGTCTTTCCTCAGGCGGCAGCGCA |
| hTR-transcription, forward | T7-GGGTTGCGGAGGGTGGGCCT |
| hTR-transcription, reverse | GCATGTGTGAGCCGAGTCCTGG |
| mTR-transcription, forward | T7-ACCTAACCCTGATTTTCATTAGC |
| mTR-transcription, reverse | GGTTGTGAGAACCGAGTTCC |
| TR-reverse transcription | GCGTCTCAACTGGTGTCGTGGAGTCGGCAATTCAGT TGAGACGCGCATGTGTGAG |
| U6-reverse transcription | GCGTCTCAACTGGTGTCGTGGAGTCGGCAATTCAGT TGAGACGCAAAATATGGAA |
| hTR, forward | TTCAGGCCTTTCAGGCCGCAGGAA |
| hTR, reverse | TGGTGTCGTGGAGTCGGC |
| mTR, forward | ACCTAACCCTGATTTTCATTAGC |
| mTR, reverse | GGTTGTGAGAACCGAGTTCC |
| U6, forward | CGCAAGGATGACACGCAAATTCG |
| U6, reverse | TGGTGTCGTGGAGTCGGC |
| MEN1, forward (protein purification) | CG GGATCC TG ATGGGGCTGAAGGCCGCCCAG |
| MEN1, reverse (protein purification) | TAGAATTCTTAGTGATGGTGATGGTGATGGAGGCCTTTGCGCTG |
| TERT promoter seq, ChIP assay, forward | 5′-ACTTTGGTTGCCCAATGC-3′ |
| TERT promoter seq, ChIP assay, reverse | 5′-AAGGAAAGGTCGGCAGGT-3′ |
| hTR, forward (For Fig. 7) | GCGAAGAGTTGGGCTCTGTCA |
| hTR, reverse (For Fig. 7) | TTCCTCTTCCTGCGGCCTGAAA |

**Table S6. Fyn CRISPR sequence**

| Santa Cruz Biotechnology Cat. No | Sequence |
|---|---|
| sc-400152-NIC Fyn Double Nickase Plasmid (h) - A: | AACAACTTCCACGCAGCCGG |
| sc-400152-NIC Fyn Double Nickase Plasmid (h) - B: | ATGGAGGTCACACCGAAGCT |

### Table S7. TR FISH probe set

| Probe name | Sequence |
|---|---|
| FLAP-Y-Cy3 | /5Cy3/AATGCATGTCGACGAGGTCCGAGTGTAA/3Cy3Sp/ |
| TRProbe1 | GCATGTGTGAGCCGAGTCCTGGGTGCTTACACTCGGACCTCGTCGACATGCATT |
| TRProbe2 | CGCGCGGGGACTCGCTCCGTTCCTTACACTCGGACCTCGTCGACATGCATT |
| TRProbe3 | TTCCTGCGGCCTGAAAGGCCTGAACTTACACTCGGACCTCGTCGACATGCATT |
| TRProbe4 | GGGCCAGCAGCTGACATTTTTTGTTTGCTTTACACTCGGACCTCGTCGACATGCATT |
| TRProbe5 | GGCTTTTCCGCCCGCTGAAAGTCAGCTTACACTCGGACCTCGTCGACATGCATT |
| TRProbe6 | GTCCCACAGCTCAGGGAATCGCGCTTACACTCGGACCTCGTCGACATGCATT |
| TRProbe7 | GCCCAACTCTTCGCGGTGGCAGTGTTACACTCGGACCTCGTCGACATGCATT |
| TRProbe8 | GCGGCCTCCAGGCGGGGTTCGGGTTACACTCGGACCTCGTCGACATGCATT |
| TRProbe9 | CCGCAGGTCCCCGGGAGGGGCGATTACACTCGGACCTCGTCGACATGCATT |
| TRProbe10 | AGAATGAACGGTGGAAGGCGGCAGGCCTTACACTCGGACCTCGTCGACATGCATT |
| TRProbe11 | CGCCTACGCCCTTCTCAGTTAGGGTTTTACACTCGGACCTCGTCGACATGCATT |
| TRProbe12 | CCCCGAGAGACCCGCGGCTGACATTACACTCGGACCTCGTCGACATGCATT |
| TRProbe13 | CCTCCGGAGAAGCCCCGGGCCGATTACACTCGGACCTCGTCGACATGCATT |
| TRProbe14 | GAAAAACAGCGCGCGGGGAGCAAAAGCACTTACACTCGGACCTCGTCGACATGCATT |
| TRProbe15 | ACAAAAAATGGCCACCACCCCTCCCTTACACTCGGACCTCGTCGACATGCATT |

**Table S8. Material list**

| Material name | Cat. no | Company |
|---|---|---|
| Mitomycin C | 11435 | Cayman Chemicals |
| Knockout™ DMEM | 10829018 | Gibco |
| Knockout™ Serum replacement | 10828028 | Gibco |
| Leukemia Inhibitory Factor | L4501 | GenDepot |
| Alkaline Phosphatase Detection Kit | SCR004 | MilliporeSigma |
| ES-E14TG2a | ATCC® CRL-1821™ | ATCC |
| DMEM, high glucose, pyruvate | 11995065 | Gibco |
| Stasis™ Stem Cell Fetal Bovine Serum | 100-125 | GeminiBio |
| Gibco™ 2-Mercaptoethanol | 21985023 | Gibco |
| L-glutamine | CA009 | GenDepot |
| Non-essential Amino acid | CA012-010 | GenDepot |
| Sh-Fyn pLKO.1 lentiviral vector | TRCN0000023381 | UMN genome center |
| Sh-Menin pLKO.1 lentiviral vector | TRCN0000040138 | UMN genome center |
| psPAX2 | 12260 | Addgene |
| pMD2.G | 12259 | Addgene |
| iMFectin Poly DNA Transfection Reagent | I7200 | GenDepot |
| Polybrene transfection Reagent | TR-1003 | GeminiBio |
| Puromycin dihydrochloride | P8833 | MilliporeSigma |
| TRAPEZE® RT Telomerase Detection Kit | S7710 | MilliporeSigma |
| Colcemid | 15364 | Cayman Chemicals |
| Fluoro-Gel II with Dapi | 17985-50 | Electron Microscopy Sci. |
| TelC-Alexa488 | F1004 | PNA Bio Inc |
| TelG-Cy3 | F1006 | PNA Bio Inc |

| | | |
|---|---|---|
| CENPB-Cy3 | F3002 | PNA Bio Inc |
| Hoechst 33258 | 16756 | Cayman Chemicals |
| RNase A | T3018L | New England BioLabs Inc. |
| Blocking reagent | 11096176001 | MilliporeSigma |
| DNAzol® reagent | 10503027 | Invitrogen |
| SYBR™ Green PCR Master Mix | 4344463 | Applied Biosystem |
| Protein G-Agarose | P9202 | GenDepot |
| NE-PER™ Nuclear and Cytoplasmic Extraction Reagents | 78835 | ThermoFisher Sci. |
| DAB Substrate | 11718096001 | MilliporeSigma |
| Recombinant Human Menin Protein | H00004221-P01 | Novus Biologicals |
| Active Fyn protein | 14-441 | MilliporeSigma |
| Recombinant Human beta Catenin protein | ab63175 | Abcam |
| BL21(DE3)-pLysS cells | L1195 | Promega |
| pGEX-5X1 | 28-9545-53 | MilliporeSigma |
| Lysozyme | 50-488-779 | Research Product Int. |
| Ni-NTA agarose | R90110 | ThermoFisher Sci. |
| Men1-pDNR-Dual | HsCD00000838 | DNASU |
| pcDNA3-HA-Sumo1 | 21154 | Addgene |
| QuikChange II Site-directed mutagenesis kit | 200521 | Agilent |
| EpiQuik™ In Vivo Universal Protein Sumoylation Assay Kit | P-8001 | EPIGENTEK |
| One-Day Chromatin Immunoprecipitation Kit | (Magna ChIP G,17-611 | MilliporeSigma |

| | | |
|---|---|---|
| Biotin-16-UTP | 11388908910 | MilliporeSigma |
| TranscriptAid T7 High Yield Transcription Kit | K0441 | Thermo Fisher Sci. |
| Streptavidin Agarose | 69203 | MilliporeSigma |
| SureStart Taq DNA Polymerase | 600282 | Agilent |
| 3-(4-chlorophenyl)-1-(1,1-dimethylethyl)-1H-pyrazolo[3,4-d]pyrimidin-4-amine (PP2) | 13198 | Cayman Chemicals |
| Dyskeratosis Fibroblast from Skin | AG04646 | Coriell Institute |
| Fyn Double Nickase Plasmid, human | sc-400152-NIC | Santa Cruz Biotechnology |
| Control Double Nickase Plasmid | sc-437281 | Santa Cruz Biotechnology |
| STEMCCA Constitutive Polycistronic (OKSM) Lentivirus Reprogramming Kit | SCR510 | MilliporeSigma |
| Recombinant Human Fibroblast Growth Factor-basic | PHG0266 | Gibco |
| TeSR™-E8™ | 05990 | StemCell Technologies |
| Matrigel® hESC-qualified Matrix | 354277 | Corning |
| Y-27632 | 72304 | StemCell Technologies |
| G418, Geneticin | 10131035 | Thermo Fisher Sci. |
| Xpert Protease Inhibitor Cocktail Solution (100X) | P3100 | GenDEPOT |
| RNase Inhibitor, Murine | M0314L | New England Biolabs. |
| N-Ethylmaleimide | E3876-5G | MilliporeSigma |

| | | |
|---|---|---|
| TransIT-X2 | MIR 6004 | Mirus |
| Lambda Protein Phosphatase | P0753S | New England Biolabs |
| YOYO®-1 | Y3601 | Invitrogen |
| Dextran Sulphate | 42867 | MilliporeSigma |
| Ribonucleoside Vanadyl Complex (VRC) | S1402S | New England Biolabs |
| NEBuffer 3 | B7003S | New England Biolabs |
| Bovine Serum Albumin | B9000S | New England Biolabs |
| tRNA | 10109517001 | MilliporeSigma |
| Xpert Prestained Protein Marker | P8502-050 | GenDEPOT |
| Xpert 2 Prestained Protein Marker | P8503-050 | GenDEPOT |

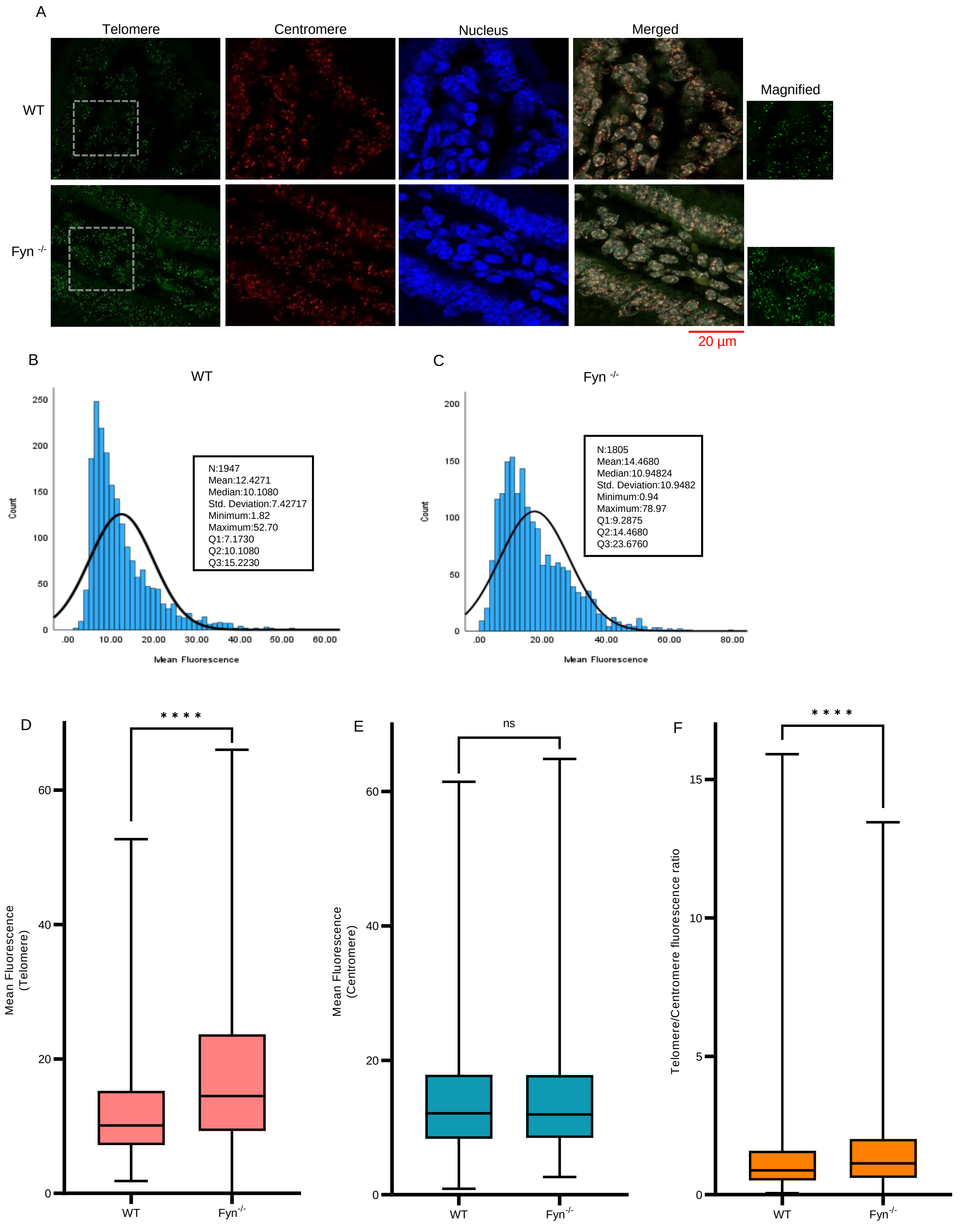


Supplementary Fig. 1

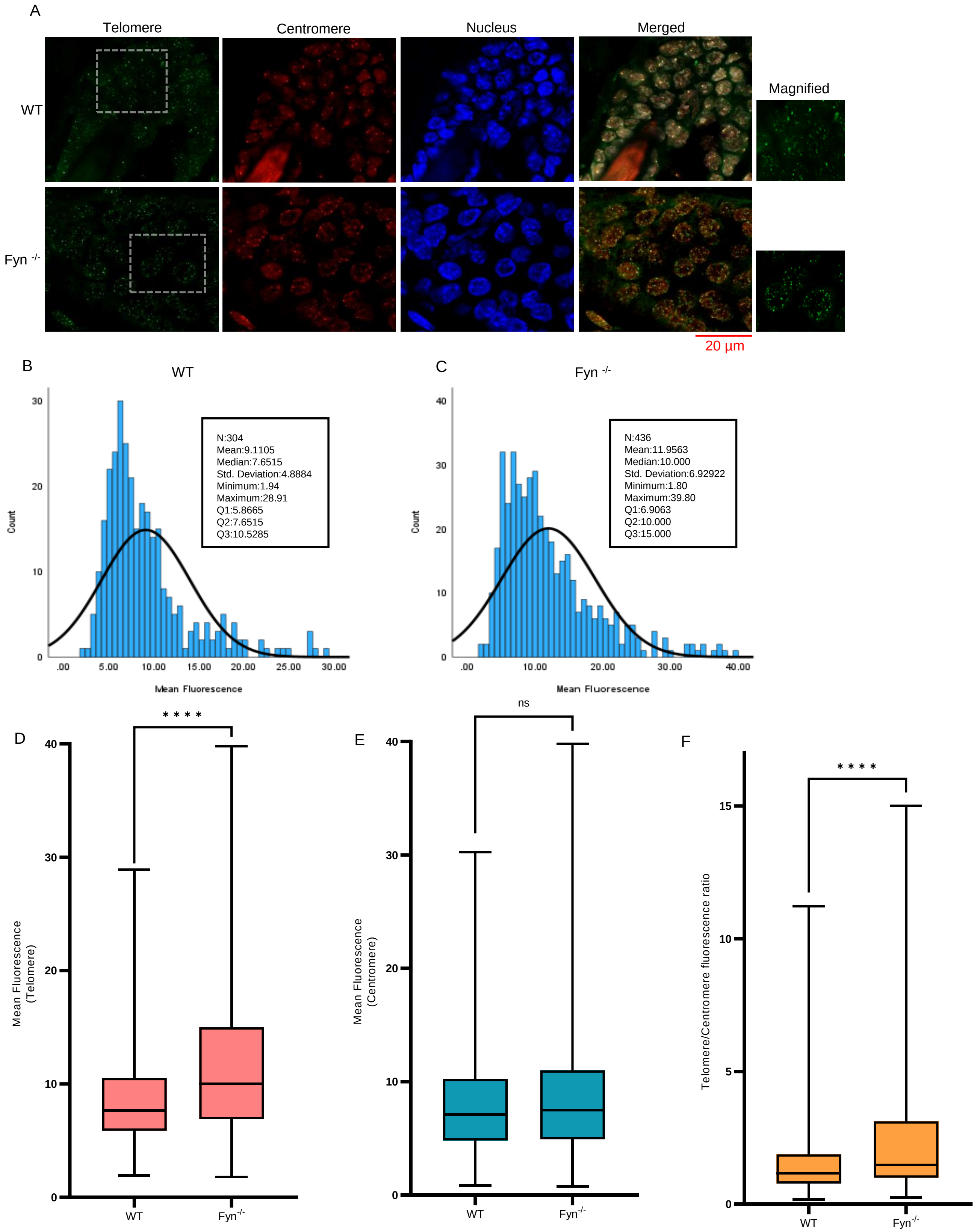


Supplementary Fig. 2

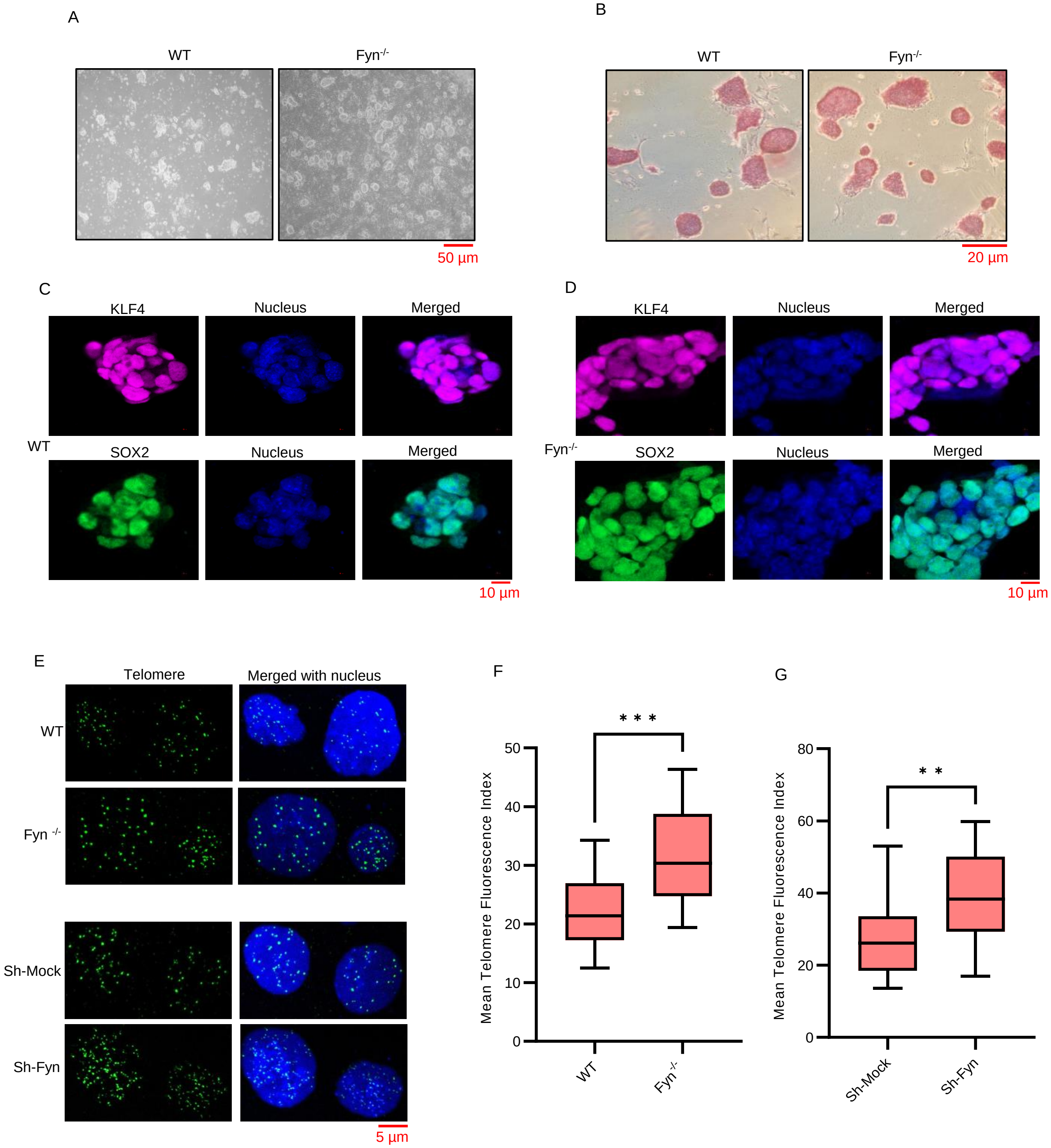


Supplementary Fig. 3

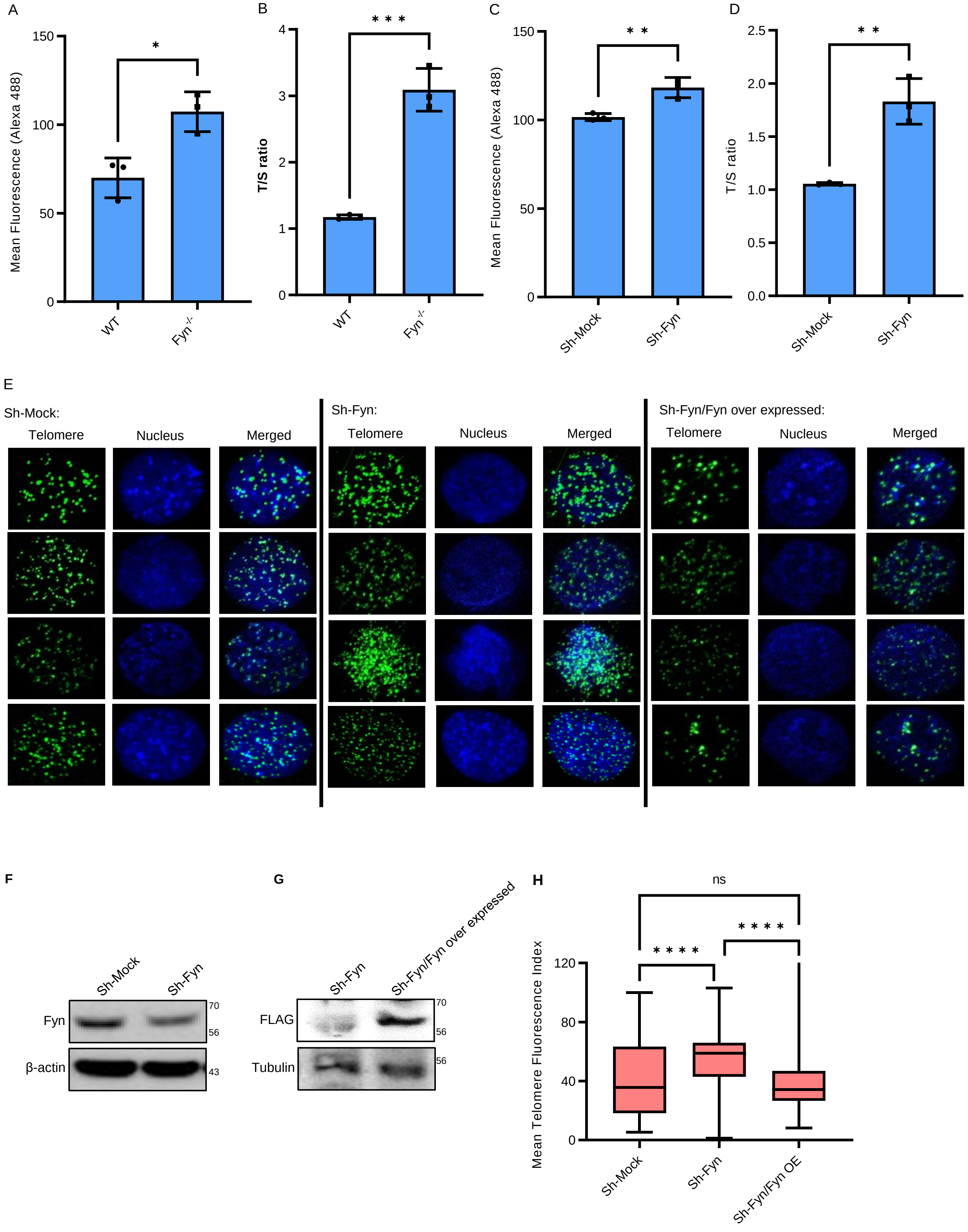


Supplementary Fig. 4

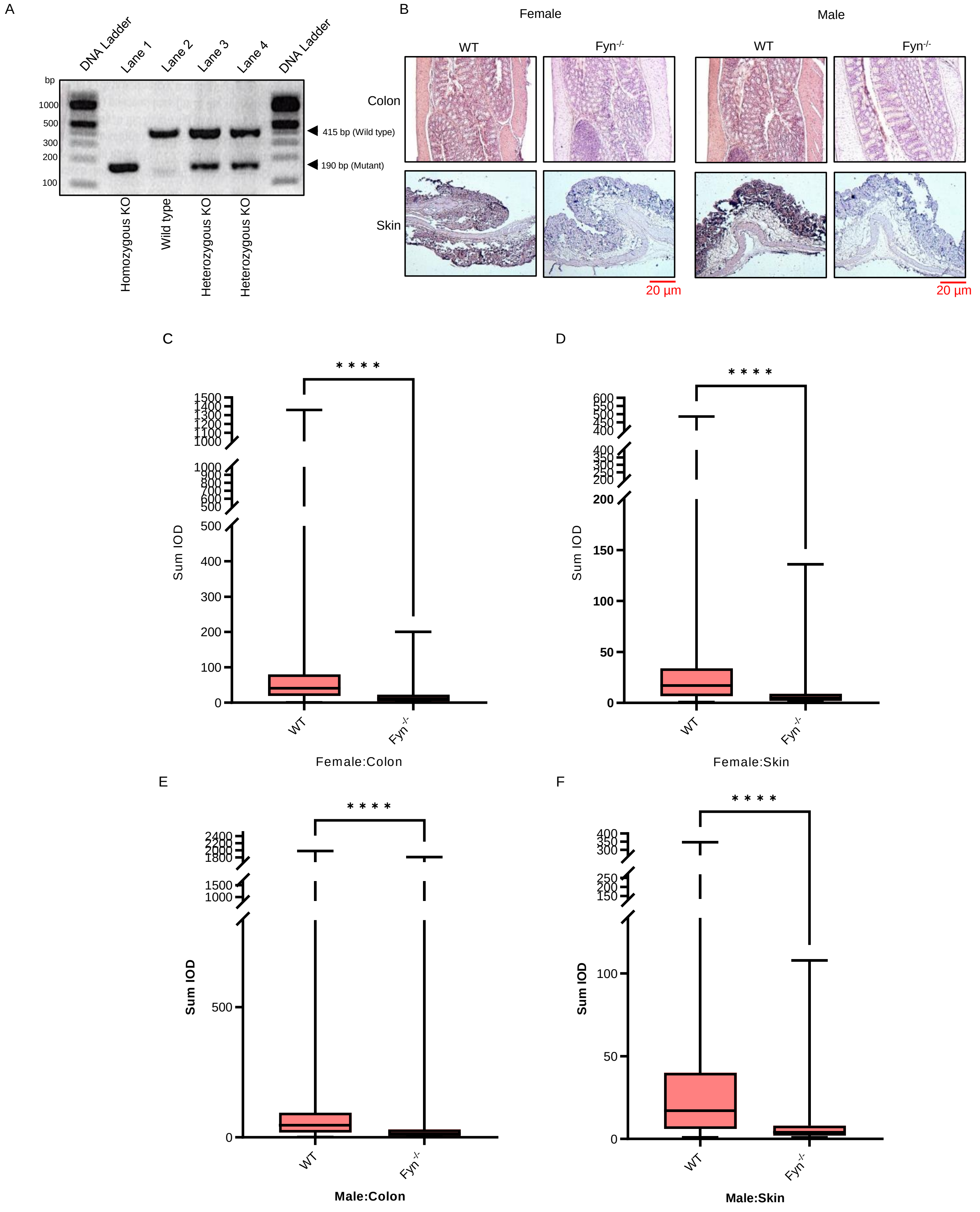


Supplementary Fig. 5

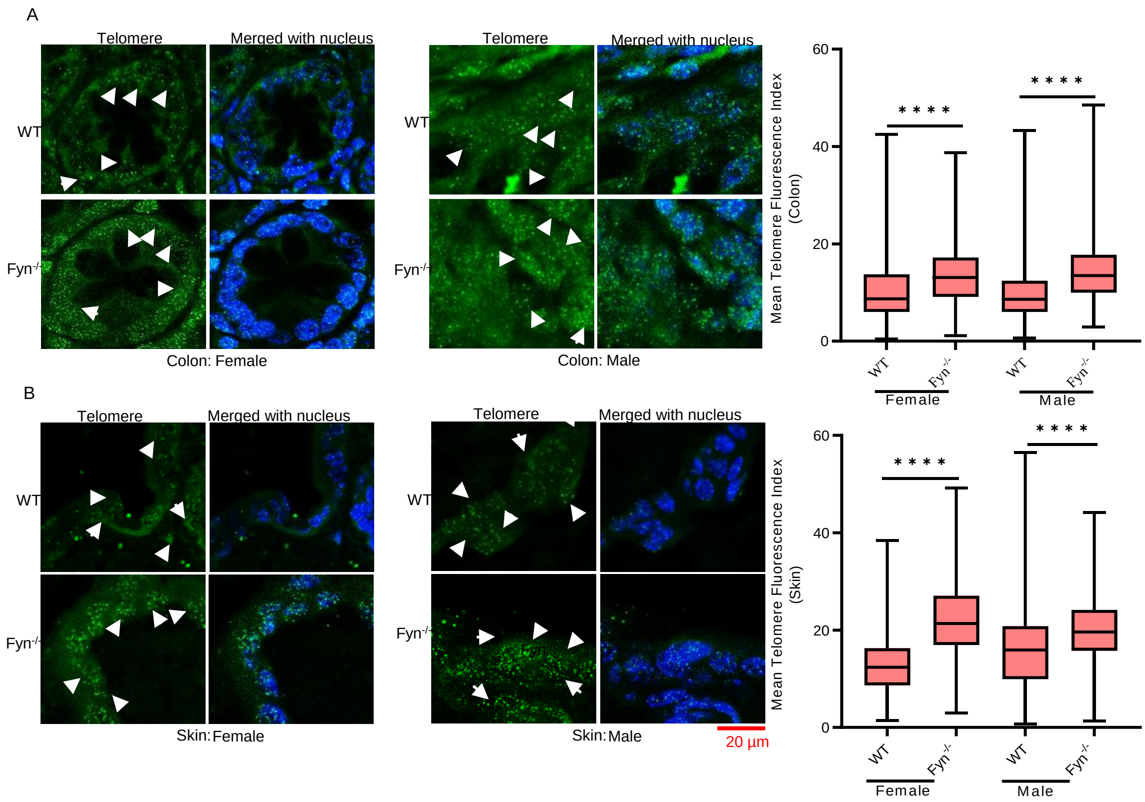


Supplementary Fig. 6

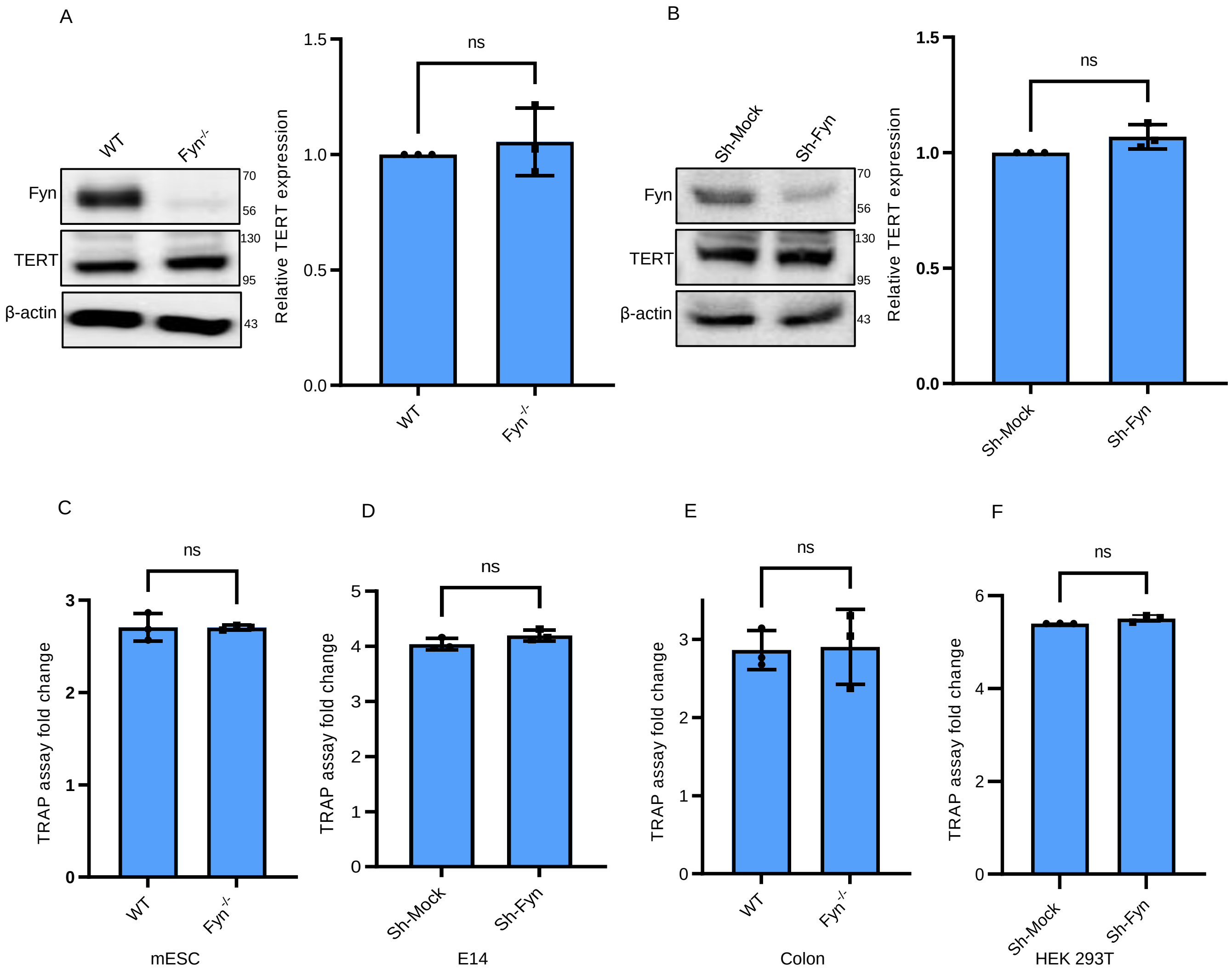


Supplementary Fig. 7

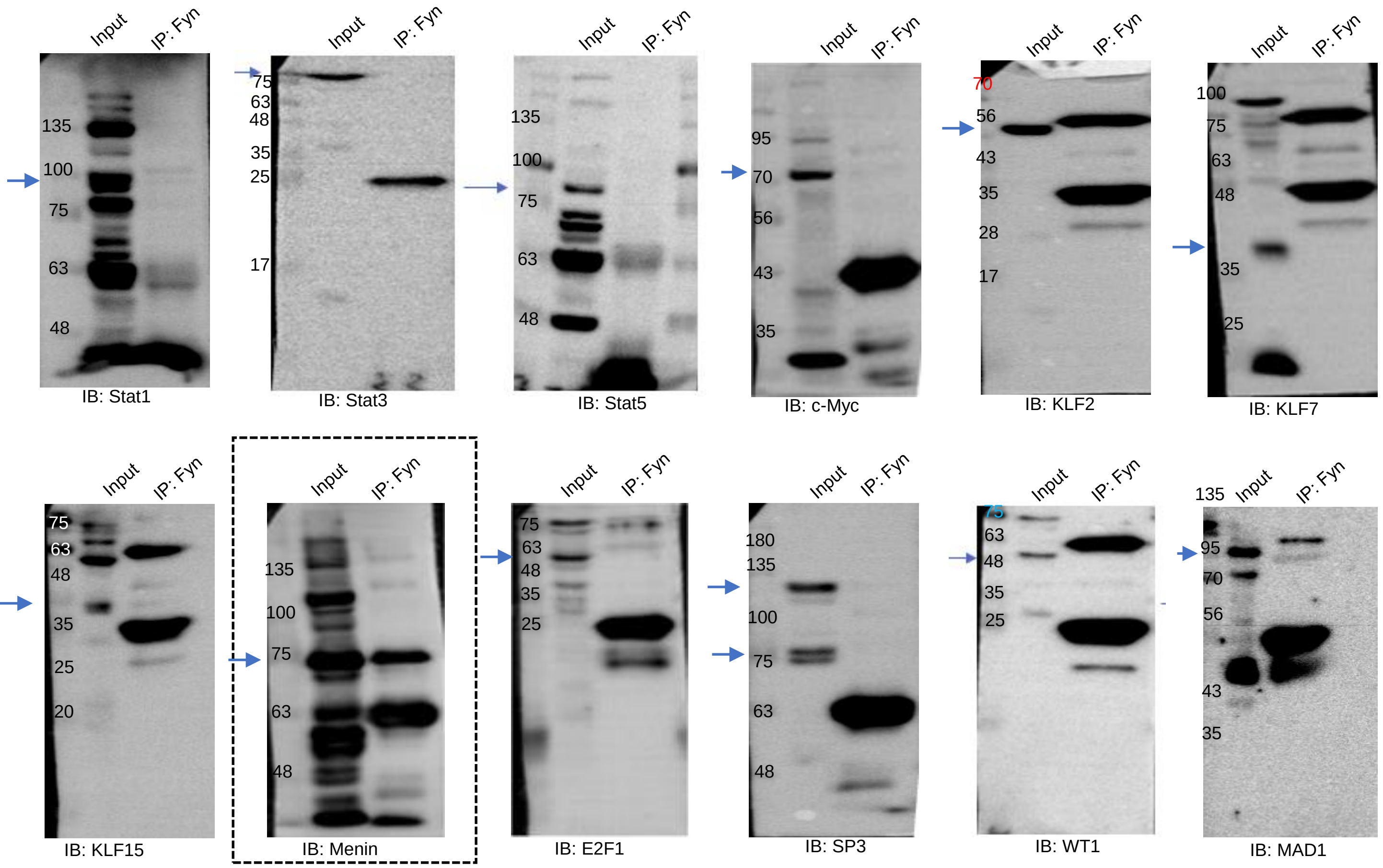


Supplementary Fig. 8

A

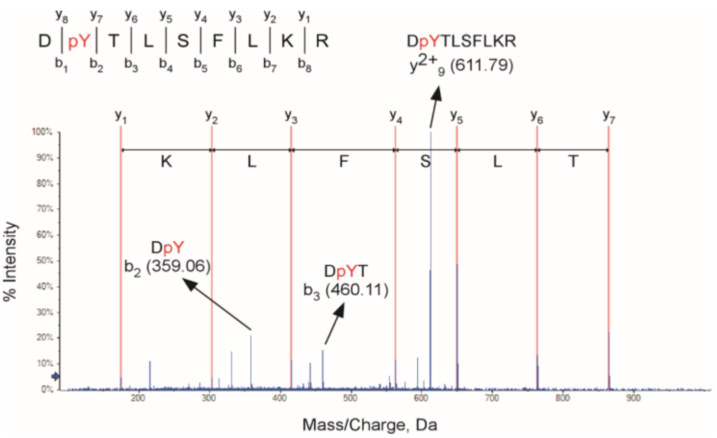


B

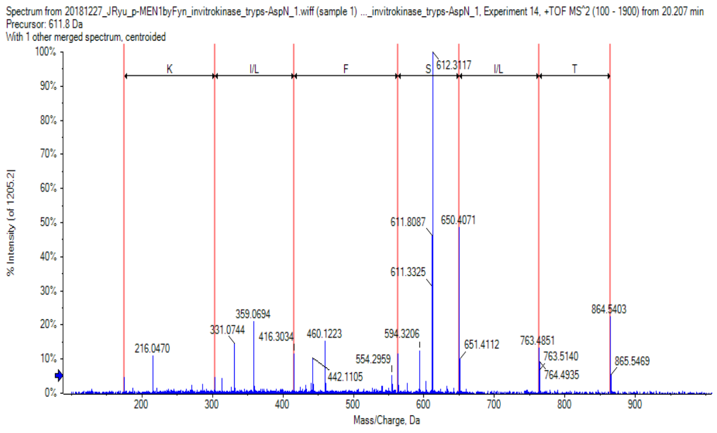


| Conf.[1] | Sc[2] | Prec m/z[3] | z[4] | Sequence | Theor m/z[5] | ΔMass[6] | Site |
|---|---|---|---|---|---|---|---|
| 99 | 12 | 611.7975 | 2 | DpYTLSFLKR | 611.7970 | 0.0009 | Y603 |

1, the confidence for the peptide identification; 2, the score for the peptide; 3, precursor mass/charge; 4, the charge for the fragmented ion; 5, theoretical precursor mass/charge; 6, the difference between theoretical molecular weight and experimental molecular weight of the matching peptide sequence

C

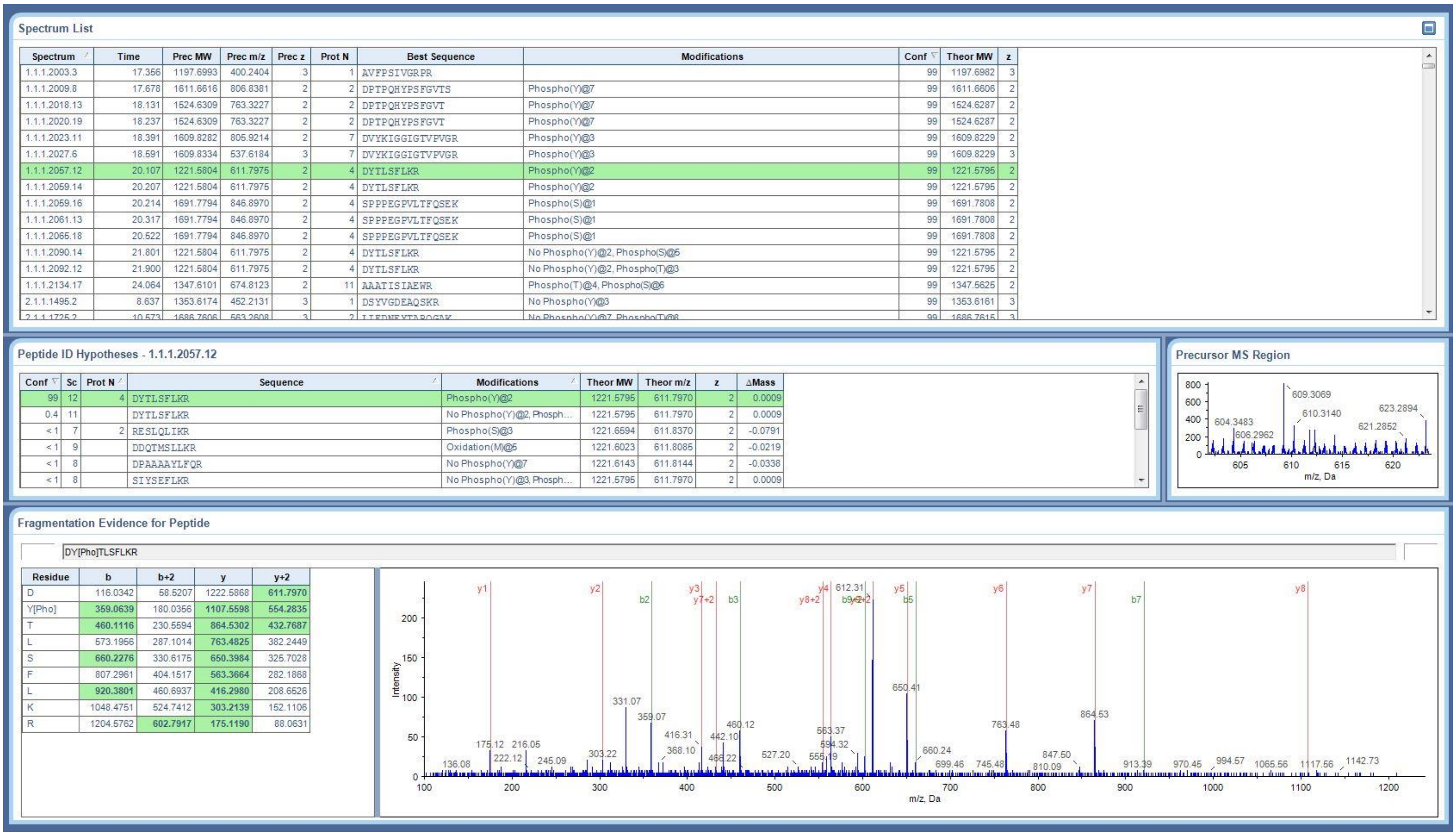


Supplementary Fig. 9

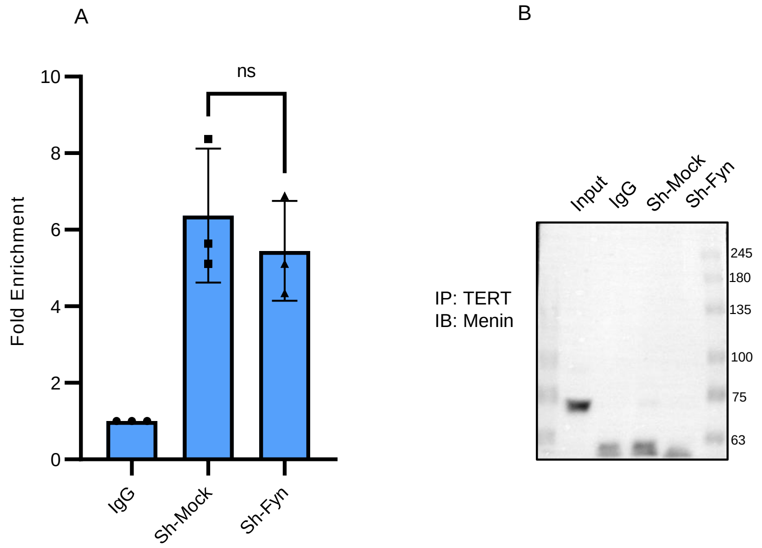


Supplementary Fig. 10

A

| Position | Peptide | Score | Cutoff | P-Value |
|---|---|---|---|---|
| 156 | GWSPVGTKLDSSGVA | 2.919 | 2.592 | 0.26 |
| 238 | SYMRCDRKMEVAFMV | 2.847 | 2.592 | 0.182 |
| 427 | RFYDGICKWEEGSPT | 2.608 | 2.592 | 0.398 |
| 493 | RGPRRESKPEEPPPP | 4.31 | 2.592 | 0.053 |
| 609 | DYTLSFLKRQRKGL* | 2.697 | 2.592 | 0.052 |
| 613 | SFLKRQRKGL***** | 3.328 | 2.592 | 0.056 |

B

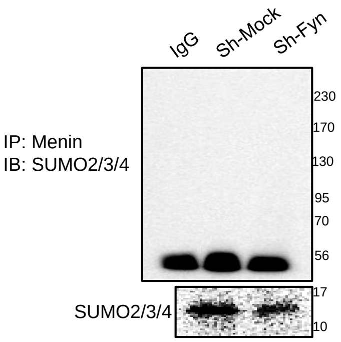


C

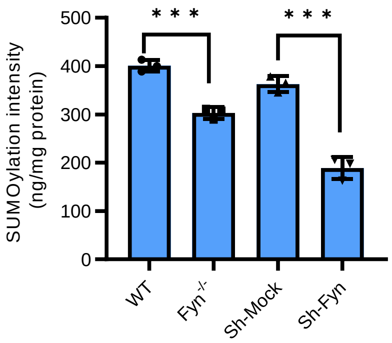


Supplementary Fig. 11

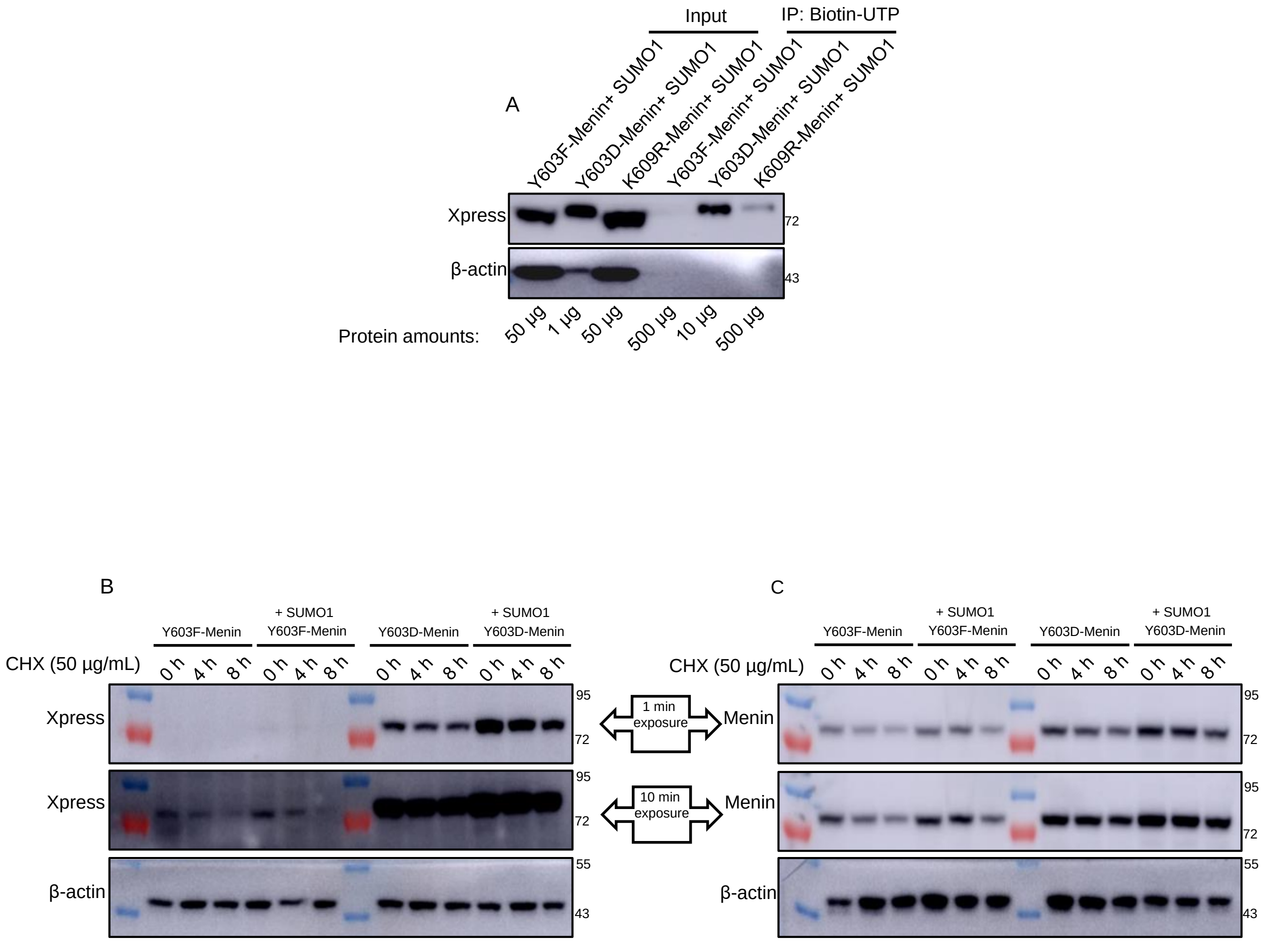


Supplementary Fig. 12

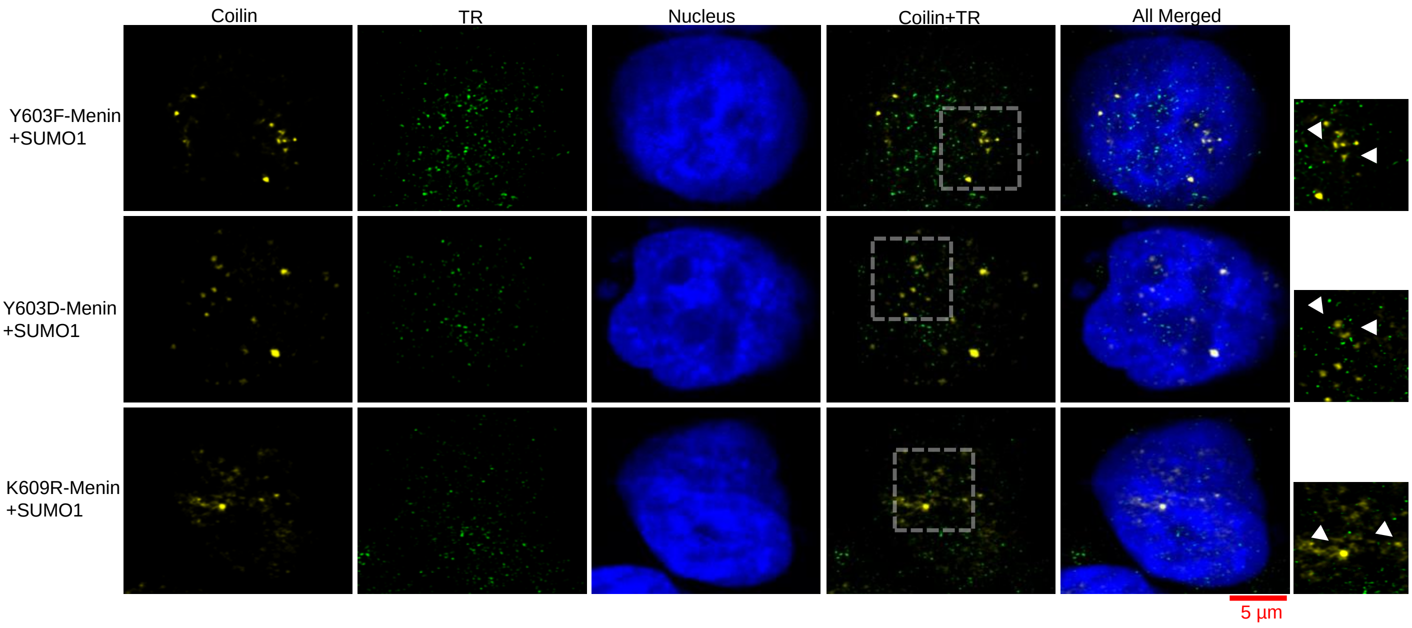


Supplementary Fig. 13

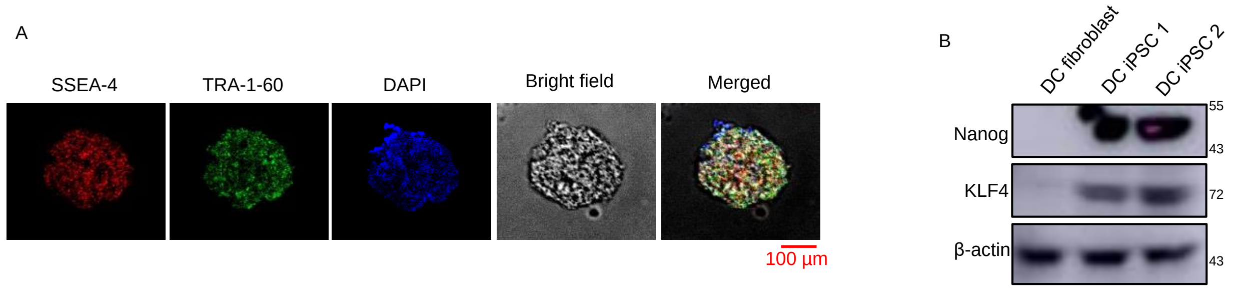


Supplementary Fig. 14
